# Updated phylogeny and protein structure predictions revise the hypothesis on the origin of MADS-box transcription factors in land plants

**DOI:** 10.1101/2023.01.10.523452

**Authors:** Yichun Qiu, Zhen Li, Dirk Walther, Claudia Köhler

## Abstract

MADS-box transcription factors (TFs) are broadly present in eukaryotes. Varying by domain architecture, MADS-box TFs in land plants are categorized into Type I (M-type) and Type II (MIKC-type). For about twenty years, Type I and II genes were considered orthologous to the SRF and MEF2 genes in animals, respectively, presumably originating from a duplication before the divergence of eukaryotes. Here, we exploited the increasing eukaryotic MADS-box sequences and reassessed their evolution. While supporting the ancient duplication giving rise to SRF- and MEF2-types, we found that Type I and II genes originated from the MEF2-type genes through another duplication in the most recent common ancestor (MRCA) of land plants. Protein structures predicted by AlphaFold2 and OmegaFold support our phylogenetic analyses, with plant Type I and II TFs resembling the MEF2-type structure, rather than SRFs. We hypothesize that the ancestral SRF-type TFs got lost in the MRCA of Archaeplastida (the kingdom Plantae *sensu lato*). The retained MEF2-type TFs acquired a Keratin-like domain and became MIKC-type upon the evolution of Streptophyta. Subsequently in the MRCA of land plants, M-type TFs evolved from a duplicated MIKC-type precursor through loss of the Keratin-like domain, leading to the Type I clade. Both Type I and II TFs largely expanded and functionally differentiated in concert with the increasing complexity of land plant body architecture. We attribute the adaptation to the terrestrial environment partly to the divergence among MEF2-type MADS-box genes and the repetitive recruitment of these originally stress-responsive TFs into developmental programs, especially those underlying reproduction.

## Introduction

MADS-box genes are a famous and intriguing gene family, broadly present in eukaryotic genomes. They encode transcription factors (TFs) which regulate diverse and important biological functions as reported in animals, fungi, plants, and protists (1). The name derives from the four founding members, Minichromosome maintenance 1 (<u>M</u> cm1) from *Saccharomyces cerevisiae*, <u>A</u> GAMOUS from *Arabidopsis thaliana*, <u>D</u> EFICIENS from *Antirrhinum majus*, and Serum response factor (<u>S</u>RF) from *Homo sapiens* (2). Animal genomes generally have two types of MADS-box genes, SRF and myocyte enhancer factor-2 (MEF2) genes that are present in one to a few copies. The budding yeast *Saccharomyces cerevisiae* has four MADS-box TFs; Mcm1 and Arg80 are related to the animal SRF, and Rlm1 and Smp1 are related to MEF2. Several phylogenetic analyses inferred the origin of SRF and MEF2 types through an ancient gene duplication event before the divergence of eukaryotes (Fig.1a,b) (3, 4, 5). The two types can be distinguished by the domains downstream of the MADS domain; while SRF type TFs are characterized by a SAM domain (SRF, ARG80 and MCM1), the corresponding region in MEF2 type TFs is referred to as the MEF2 domain (6). The crystal structures of several MADS-box TFs have been resolved, including human SRF and budding yeast MCM1, human MEF2A and mouse MEF2C. The conserved MADS domain comprises an alpha helix, followed by two antiparallel beta strands. After which, the MEF2-type and SRF-type structures differ in the second alpha helix, constituted by the SAM or MEF2 domain, respectively, where a kink in the SAM domain of SRF-type TFs changes the orientation of the second helix in the opposite direction to that of MEF2-type TFs (Fig.2a). The two alpha helices and the connecting beta strands build up the interface for TF dimerization and DNA-binding.

In land plants, MADS-box TFs (also referred to as AGAMOUS-like (AGLs) in the model plant *Arabidopsis thaliana*) have evolved to be a flourishing family with typically as many as 50 to over 100 members in angiosperm species, in sharp contrast to the much smaller family sizes of MADS-box TFs in other eukaryotes (7). According to the domain architecture, the MADS-box TFs in land plants can be specified into two types: the Type I TFs are usually referred to as M-type, since they share no well-characterized conserved domain following the MADS domain, while the Type II TFs typically have the <u>M</u>ADS domain followed by the I ntervening, <u>K</u> eratin-like and <u>C</u> −terminal domains, so they are also known as MIKC-type. The I domain is analogous to the SAM or MEF2 domain in animal MADS-box TFs in terms of location and function, and the K domain is likely specific for plants (4, 8). Due to their critical roles in establishing floral organ identity in angiosperms, the Type II MADS-box genes have been studied extensively. In contrast, Type I MADS-box genes have only been identified along with the first angiosperm genome of *Arabidopsis thaliana*, and gradually emerging studies have linked their functions to the development of the female gametophyte and endosperm (9, 10, 11). Besides the distinct domain arrangements of their encoded proteins, land plants Type I and II MADS-box genes vary in expression patterns, numbers of exons, and noticeably, compared with Type II MADS-box genes, Type I genes have evolved faster and more frequently undergone duplication and loss (8, 12).

Among many known regulatory functions, MADS-box TFs act as major regulators of plant reproduction and have been closely connected with the rise of flowering plants to ecological dominance (13, 14). The family size of MADS-box genes has been linked to the complexity of the plant body plan (3, 14, 15). Thus, identifying the origin and resolving the subsequent diversification of plant MADS-box genes is required to understand the evolutionary success of land plants.

Upon the discovery of Type I MADS-box genes, a timely survey suggested that Type I and II genes in plants are orthologous to the SRF and MEF2 genes in animals, respectively (4). Based on this model, an ancient duplication before the divergence of the extant eukaryotic lineages gave birth to the two classes of MADS-box genes in plants, likewise in animals and other eukaryotes (Fig.1). This model has been influential in the field of MADS-box evolution and served as a basis for investigations of MADS-box gene evolution across all phylogenetic scales. Nevertheless, some thoughtful critique on this model has been neglected for a while (14, 16). As already noted by the authors of the model (4), the clustering of Type I TFs in plants and SRF-type TFs in animals and fungi was not well supported in the original study based on only a few Arabidopsis, animal, and fungal sequences.

**Fig.1.**
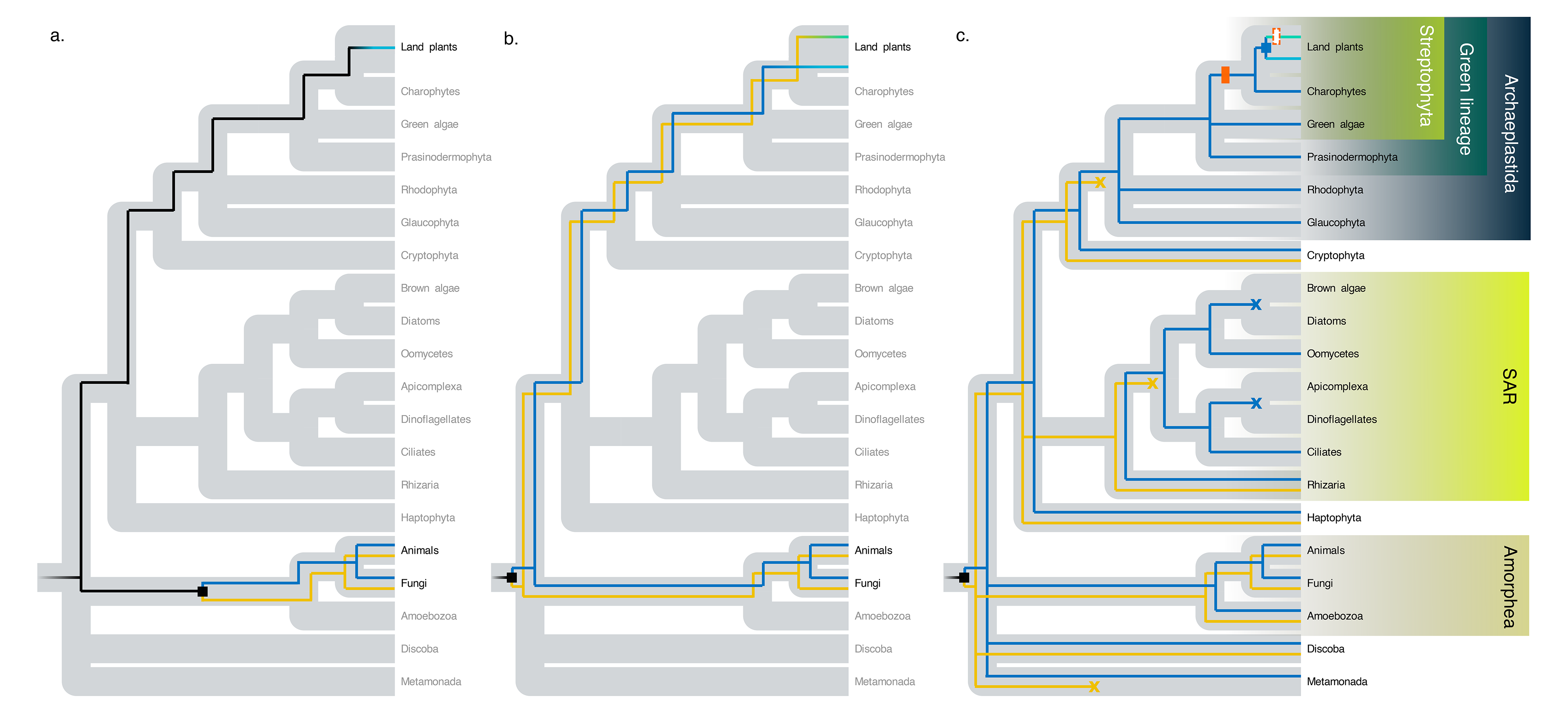
The models of MADS-box transcription factor (TF) evolution. **a.** The model by Theissen et al. (3): Before the identification of M-type MADS-box TFs, known MADS-box TFs in plants were all belonging to the MIKC-type. The two types of MADS-box TFs in animals and fungi, SRF and MEF2, were considered duplicates that diverged after the split of the plant and animal lineages. **b.** The model by Alvarez-Buylla et al. (4): Upon the identification of M-type MADS-box TFs in the genome of *Arabidopsis thaliana*, they were named Type I MADS-box TFs and considered orthologous to SRF-type TFs in animals and fungi, while MIKC-type in plants were grouped as Type II and clustered with MEF2-type in animals and fungi. Type I and II MADS-box genes originated by a hypothesized ancient duplication event predating the plant and animal divergence. **c.** The model proposed in this study: With a broad survey across eukaryotes, the ancient duplication giving rise to SRF- and MEF2-type can be inferred as early as the origin of the most common recent ancestor of all living eukaryotic groups (consistent with model b). In several modern lineages both types have been retained; however, in the lineage of Archaeplastida, SRF-type TFs got lost, and only MEF2-type TFs continued evolving. Subsequently, Type I and II MADS-box TFs originated by duplication of a MEF2-type precursor in the most recent common ancestor of all land plants. Black, primitive MADS- box TFs; yellow, SRF-type; blue, MEF2-type; green, Type II; cyan, Type I. Square, gene duplication event; cross, gene loss. Orange bar, gain of the K domain; empty orange bar, loss of the K domain.

The burst of available genome sequences of plants as well as other major clades of eukaryotes encouraged us to revisit the origin of MADS-box genes in land plants. In particular the genomes of Charophytes, the paraphyletic algal relatives of land plants, have shed light on the evolution of gene families underlying the successful terrestrialization of land plants (17, 18). There, in the charophyte *Klebsormidium flaccidum*, the only present MADS-box gene belongs to Type II, coding for a MIKC-type TF (17). Similarly, in the *Chara braunii* genome, the three identified MADS-box genes belong to Type II, since they are all related to MEF2 genes (18). Furthermore, in the genomes of the green algae *Chlamydomonas reinhardtii*, *Ostreococcus tauri*, *Ostreococcus lucimarinus* and the red algae *Cyanidioschyzon merolae*, the annotated MADS-box genes are identified as MEF2-type, despite that the encoded TFs lack a K domain (14, 15). Surprisingly, SRF-type MADS-box genes have so far never been found in the charophycean, green or red algae. If Type IMADS-box genes in land plants are descendants of the ancestral SRF-type genes as stated by the original model (4), the orthologous SRF-type genes would have been lost convergently and repeatedly in all of successive sister groups to land plants, the paraphyletic algal relatives. Thus, the systematic lack of SRF-type TFs in these lineages challenges the orthology between Type I MADS-box TFs in land plants and SRF-type TFs in animals and fungi. Enabled by the newly available genome sequences and the unprecedented power to predict protein structures based on amino acid sequences, we reassessed the origin of plant Type I MADS-box genes and propose an alternative model explaining the evolution of MADS-box genes in the green lineages.

## Results

### Ancient duplication of SRF and MEF2 clades is supported using newly available genome sequences

Capitalizing on newly available genome sequences spanning the broad phylogeny of eukaryotes, we re-evaluated the original phylogenetic model of MADS-box gene evolution with extended sample sequences. We collected published genomes of different lineages of eukaryotes and identified MADS-box genes in 175 species (Supplementary table 1). The species represent seven currently accepted eukaryotic groups: 1. Archaeplastida (the kingdom Plantae *sensu lato*, e.g., streptophytes including land plants, green algae, Prasinodermophyta, red algae, and glaucophytes), 2. Cryptista, 3. Haptista, 4. SAR supergroup which are Stramenopila (e.g., brown algae, diatoms, oomycetes) / Alveolata (e.g., ciliates, dinoflagellates, Apicomplexa) / Rhizaria, 5. Amorphea (e.g., animals, fungi, amoebae), 6. Discoba, and 7. Metamonada. The latter two were previously collectively referred to as Excavata (19).

We selected protein sequences of MADS-box TFs from the diverse eukaryotic groups. To perform multiple sequence alignments (MSAs), we extracted the conventional MADS domain sequences of about 60 amino acids in length as defined in previous studies (6) extended by the corresponding regions of SAM/MEF2/I domains, so that the extended MSAs structurally cover the functional unit of two alpha helices and the connecting beta strands. We inferred phylogenetic trees with maximum likelihood, Bayesian inference, and neighbour-joining. All three methods consistently found that the MADS domain sequences naturally form two major clades, corresponding to current SRF and MEF2 lineages, as referenced by the known SRF and MEF2 sequences from animals, fungi, and amoebae (Fig.3,4; Supplementary Fig.S1,2). This finding supports the overarching hypothesis that an ancient duplication of a MADS-box gene gave rise to the SRF-type and MEF2-type precursors. We found both SRF- and MEF2-types genes in nearly all of the surveyed Amorphea species, as well as Cryptista and Haptista. Furthermore, two species in Discoba, representing a distinct group distantly diverged from the plant and animal lineages, have both types of MADS-box genes (Supplementary table 1). Together, our results provide supporting evidence for the presence of SRF- and MEF2-type MADS-box genes early before the diversification of major groups of extant eukaryotes.

### Land plant Type I and II MADS-box TFs are both MEF2-type

Previous studies concluded that Type I MADS-box genes in land plants are more closely related to the SRF genes in animals (Fig.1b) (4, 20). In contrast, our phylogenetic analyses using three different phylogenetic approaches consistently suggested that Type I MADS-box TFs in land plants clustered within the MEF2 clade, which includes plant Type II TFs (Fig.3,4; Supplementary Fig.S1,2,3). Thus, both Type I and II genes are inferred to be MEF2-type and no SRF-type gene is present in the extant land plants. In addition, we carried out approximate unbiased (AU) tests (21) to compare the two competing phylogenetic trees of MADS-box TFs. One topology represents our new phylogeny, which groups plant Type I and II both within the MEF2 clade; the other is a constraint phylogeny forcing the plant Type I clade into the SRF clade reflecting the previous model. The AU tests significantly rejected the topology with the clustering of plant Type I TFs in the SRF clade (Supplementary Fig.S4).

### MADS domains of SRF-type and MEF2-type TFs form distinct structures

The functional units of SRF- and MEF2-type (extended MADS domain) are known to form distinct protein structures (Fig.2a). We applied AlphaFold2 (22) to predict the structures of identified eukaryotic MADS-box TFs and compared their overlay patterns with the resolved SRF- or MEF2-type MADS domain structures (Fig.2b,c; Supplementary table 2). For the structures of TFs that belong to the MEF2-type inferred by phylogeny, the human SRF (HsSRF) structure was frequently not considered most similar, but rather the human MEF2A (HsMEF2A) structure. In contrast, the sequences present in the SRF clade fitted significantly better to the HsSRF structure than the HsMEF2A structure (Fig.2b,c; Supplementary Fig.S5). We also applied OmegaFold (23) to predict the structures of a larger set of MADS domain sequences (Supplementary table 3). Different from AlphaFold2 which makes use of MSAs for prediction, OmegaFold is able to predict the structures based on their individual primary sequences, without an explicit alignment. The results of structure similarities obtained from OmegaFold predictions agree with the comparisons based on the AlphaFold2 predictions (Fig.2e,f; Supplementary Fig.S5). Hence, the predicted structure of the MADS domain supports the classification of proteins into SRF- or MEF2-type.

**Fig.2.**
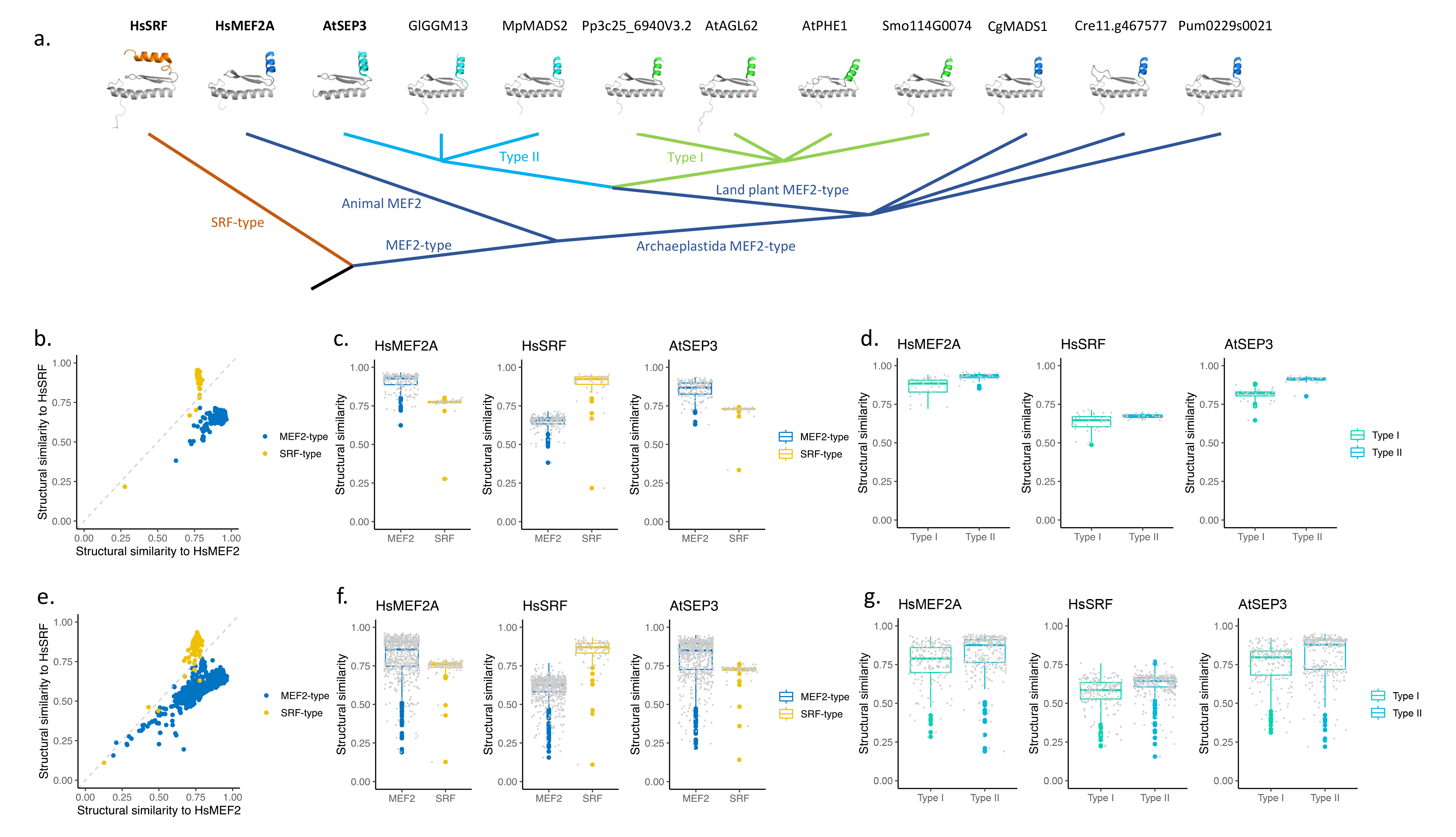
Protein analyses of MADS-box TFs. **a.** Crystal structures (bold) of human SRF and MEF2A, and Arabidopsis SEP3 that were used as templates for structural comparisons (A-chains only), and AlphaFold2-predicted structural models of Archaeplastida MADS-box proteins including Type I and II TFs in land plants. Models were trimmed to the relevant structural segments matching the template structures. Structures were drawn using Pymol (65). **b.c.** Similarity scores (TM-scores, with “1” indicating high (perfect), “0” no (random) agreement) of all predicted structural models by AlphaFold2 to human SRF and MEF2A, and Arabidopsis SEP3. SRF-type (yellow) and MEF2-type (blue) TFs were inferred by phylogeny. **d.** Similarity scores of predicted structural models by AlphaFold2 for land plant MADS-box TFs to human SRF and MEF2A, and Arabidopsis AtSEP3. **e.f.g.** Similarity scores of predicted structural models by OmegaFold.

Based on AlphaFold2 and OmegaFold predictions, Type II TFs in land plants resemble the HsMEF2A structure (Fig.2d,g). Similarly, Type I TFs in land plants all mapped better to the HsMEF2A compared to the HsSRF structure. The second helix of Type I TFs was not predicted to be twisted by a kink, as found in SRF-type TFs. There is no experimentally resolved structure for a plant Type I MADS-box TFs available thus far. The only resolved crystal structure of a plant MADS-box protein is that of the Arabidopsis Type II TF SEPALLATA 3 (AtSEP3), which, as expected, displays a MEF2 structure (24). The predicted models for both Type I and II TFs in land plants, did align well with the AtSEP3 structure (Fig.2d,g), though Type II TFs had higher structural similarity scores to AtSEP3, consistent with the fact that they are more closely related. In Type II TFs, the I domain forms a helix, resembling the second helix formed by the MEF2 domain in the Amorphea MEF2-type TFs. Correspondingly, the second helix in the predicted structures of the Type I TFs is formed by an I-like domain region (24). While initially not defined in early studies (4, 12), the I-like domain in the Arabidopsis Type I TFs has been shown to be required for both dimerization and DNA binding, functionally equivalent to the I domain in Type II TFs (24).

### Loss of SRF-type genes in the most recent common ancestor of Archaeplastida

The origin and divergence of Type I and II genes in land plants can be further inferred from the sister green lineages (Fig.3,4; Supplementary Fig.S1,2,3). In line with previous findings, the presence of only MEF2-type genes and the absence of SRF were observed in the genomes of streptophytic algae, a series of successive sister groups of land plants. The loss of SRF-type genes may be tracked back to before the diversification of the whole Archaeplastida clade. In green algae and a third lineage of green plants, Prasinodermophyta, represented by *Prasinoderma coloniale*, there is no confidently predicted SRF-type gene either. In a few species belonging to the core Chlorophyta, we found some genes harboring an open reading frame coding for a partial SRF-like MADS domain, but these SRF-like domains are quite divergent from SRF-type sequences found in other eukaryotic lineages, indicated by long branches, low support scores and inconsistent positions in our phylogenetic analyses (Supplementary Fig.S6). Their best BLASTP hits against all other MADS-box sequences were fungal SRF-type sequences, indicating that they probably arose from a horizontally transferred fragment in the most recent common ancestor (MRCA) of core chlorophytes. Likewise, in red algae, which are sister to the green plants, no SRF-type gene was identified, neither in the glaucophyte *Cyanophora paradoxa*. The lack of SRF-type genes in the Archaeplastida lineage is unlikely a consequence of multiple independent losses. Instead, it raises the more parsimonious hypothesis that the MRCA of the Archaeplastida lineage did not inherit an SRF-type gene, further supporting that only MEF2-type genes gave birth to Type I and II MADS-box genes in land plants.

**Fig.3.**
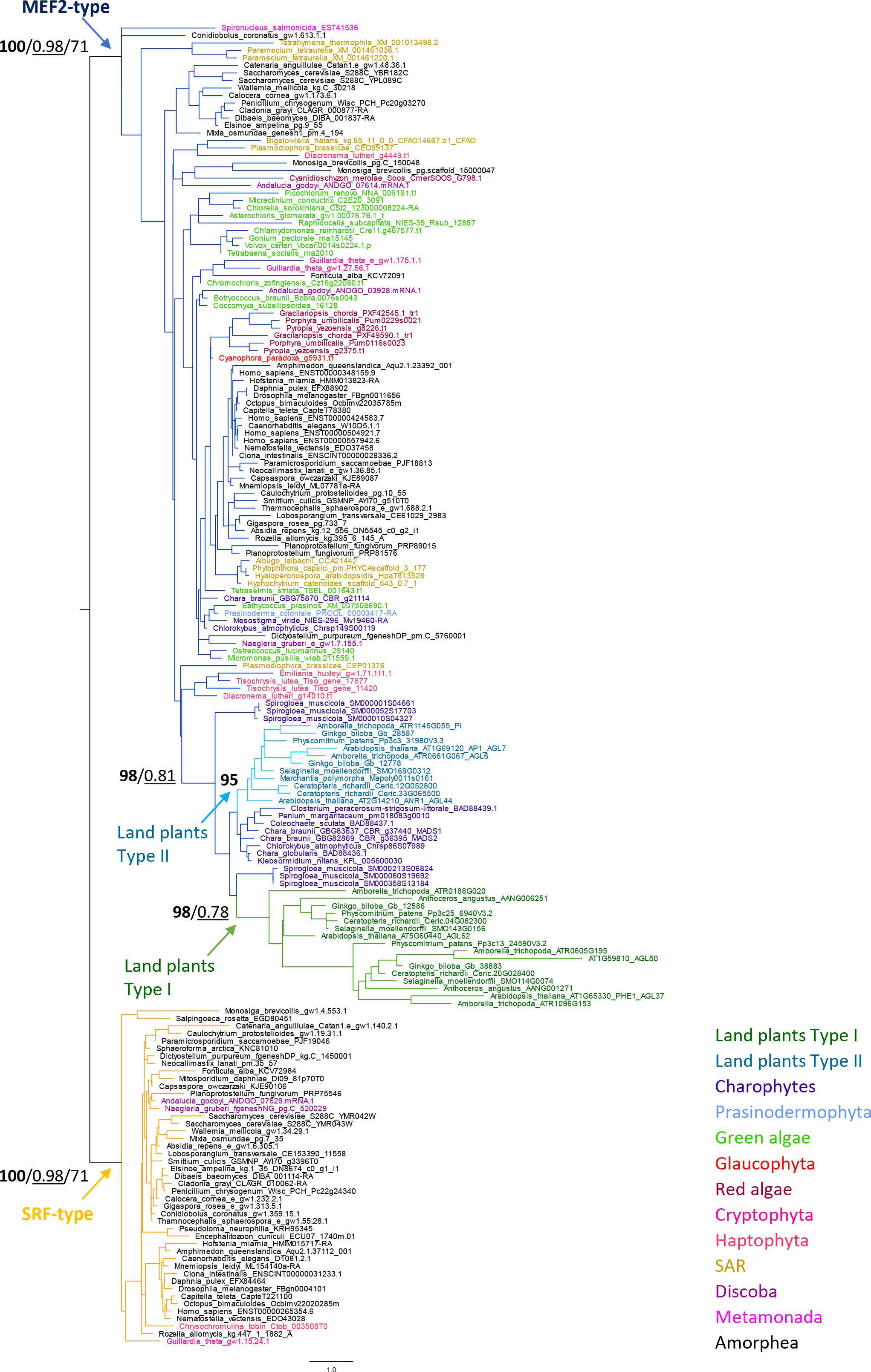
Maximum-likelihood (ML) tree of selected MADS domains (extended definition, the first and second alpha helix and the connecting beta strands). Numbers for given branches of interest are support values: bootstrap values in ML trees (bold) / posterior probability in Bayesian inference (underlined) / bootstrap values in neighbor-joining trees. Branch color: yellow, SRF-type; blue, MEF2-type; green, Type II; cyan, Type I. Sequence IDs colored by category as listed on the right.

**Fig.4.**
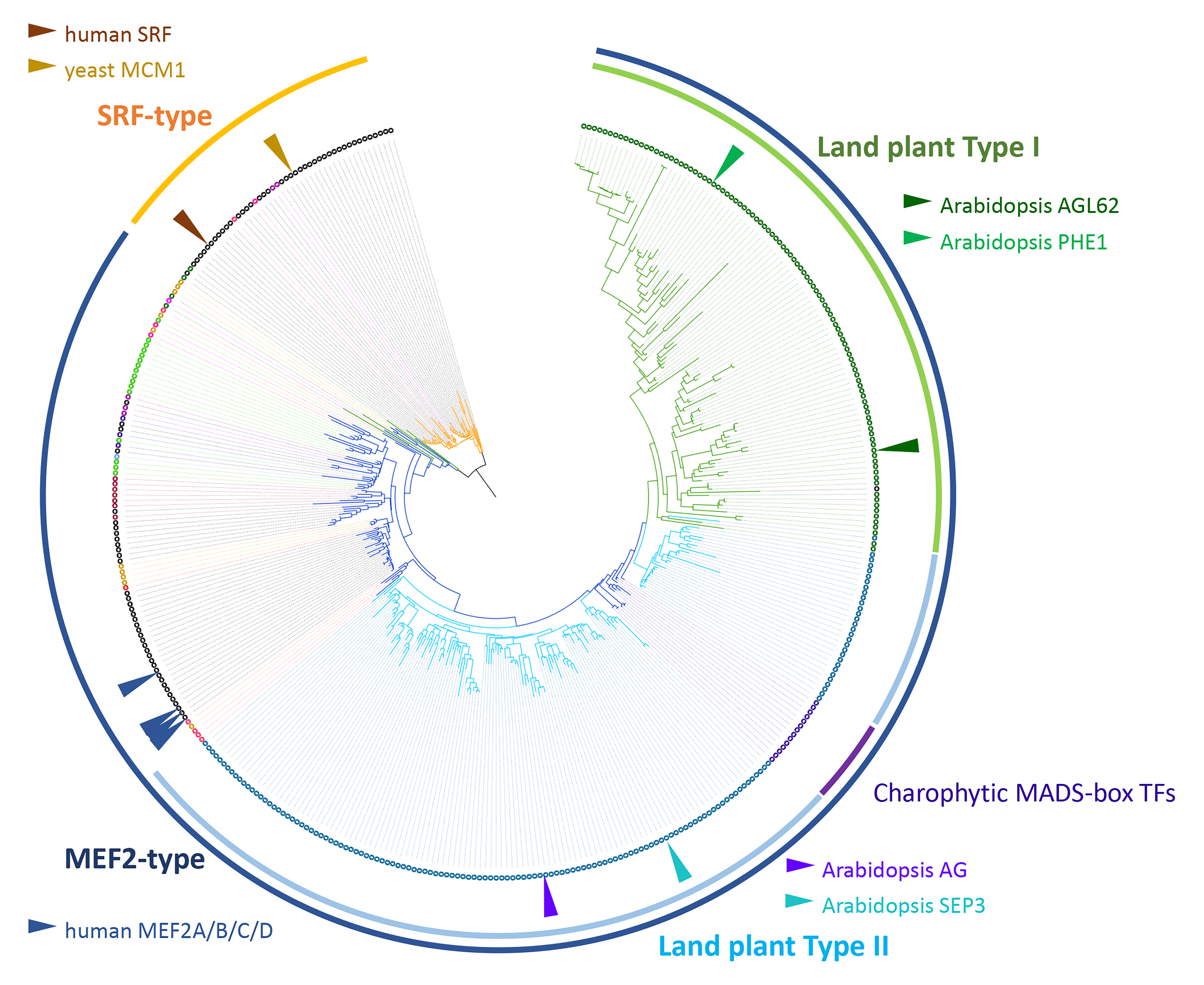
Maximum likelihood tree of extended MADS domain sequences including all MADS-box TFs from eight representative land plants. Branches and clades are colored by category as labels aside. Sequence IDs and taxon color codes are the same as the expanded tree in Fig.S2. Arrows point to several TFs with known function as the landmarks for each type.

In agreement with this evolutionary scenario, the predicted structures of Archaeplastida MADS-box TFs with or without a K domain all have higher similarity scores to the HsMEF2A and AtSEP3 structures than to the HsSRF structure (Fig.2a; Supplementary Fig.S5). All these MEF2-type structures share the Intervening or MEF2 domain-like region, which constitutes the second helix with no kink. Thus, the predicted protein structures mirror the new phylogeny (Fig.1c,2a).

### Sporadic losses of MADS-box TFs across eukaryotes

Except for Archaeplastida, some other eukaryotic lineages have only either SRF- or MEF2-type genes (Supplementary table 1). For example, three species in Microsporidia, a group of unicellular parasites closely related to fungi and *Sphaeroforma arctica* and *Thecamonas trahens*, belonging to successive sister groups of animals and fungi, they all have only SRF-type genes. In Haptista, *Chrysochromulina tobin* has only SRF-type genes, while three other species have only MEF2-type genes, suggesting reciprocal losses after the divergence from the MRCA comprising both types. Some SAR species, like ciliates, oomycetes and the cercozoan *Plasmodiophora brassicae* have only MEF2-type genes. Some surveyed species belonging to the green algae, the brown algae, diatoms, dinoflagellates and several Metamonada and Discoba protists among others have no extant MADS-box genes. To rule out the possibility that the observed absence of a certain type of MADS-box genes is a result of incomplete gene annotations, we scanned these genomes with the profile hidden Markov model for known MADS domains (PF00319) from Pfam (25). We identified some unannotated and incomplete MADS- domains in a few species (Supplementary table 1). Importantly however, no SRF-type gene was detected in the Archaeplastida. Therefore, the proposed SRF-type gene loss in the MRCA of Archaeplastida is not an artifact due to incomplete gene prediction.

## Discussion

### A new hypothesis on the origin of Type I and II MADS-box genes in land plants

Supported by an updated phylogeny and predicted protein structures, we propose that in the land plant lineage Type I MADS-box TFs arose as a second clade of MEF2-type TFs, sister to the Type II TFs. The birth of Type I and II MADS-box genes was the result of a gene duplication event of a MEF2-type ancestral gene in the MRCA of land plants, which was followed by rounds of gene duplication largely expanding the MADS-box gene family. This new model of the MEF2-type origin is also favored by the principle of parsimony considering the absence of SRF-type genes in the sister lineages of Archaeplastida, specifically those of streptophytic algae (Fig.1c).

The major difference between our model and the previously proposed model is the origin of plant Type I MADS-box genes. Type I genes are known to have high substitution rates (12), which is reflected by long branches in previous phylogenetic analyses and ours as well (Fig.3,4). Previous studies claiming that Type I genes in Arabidopsis are closely related to fungal and animal SRF-type genes suffered from biased and inadequate sampling (4). The limitation in sequence sampling also affected another early model, which suggested that Type I genes (referred to as M-type in that study) are polyphyletic, while Type II genes might be MEF2-like genes (26). The upsurge of eukaryotic genomes has been filling the gaps between distantly related animal and plant sequences, making it possible to break the long branches and improve the phylogenetic resolution of the MADS-box gene family. Our new investigation covering diverse, previously underrepresented protist groups provides comprehensive support for the hypothesized ancient duplication of SRF- and MEF2-types before the divergence of extant eukaryotes. Besides, the Charophytic genomes serve as great references for the gene family evolution in land plants. We thus took the opportunity to revisit the evolution of the MADS-box gene family that was likely a key driver for plants adapting to land ecosystems.

The identification of Type I TFs in land plants as members of the MEF2-type lineage suggests that plant Type I and II should both be considered as "Type II" as defined by Alvarez-Buylla et al. (4). Thereby, to avoid future confusion in nomenclature, we propose that referring to “Type I” and “Type II” should be restricted to MADS-box TFs in land plants, considering that these terms have been widely used in the literature in plant sciences. To differentiate the two clades of MADS-box TFs that originated from the ancient eukaryote-wide duplication, we suggest referring to them as "SRF-type" and "MEF2-type", respectively, as “SRF” and “MEF2” have been commonly used even before the proposed categories by Alvarez-Buylla et al. (4). Additionally, we recommend to use “M-type” and “MIKC-type” only when distinguishing the MADS-box TFs in plants by domain architecture rather than their origin.

### Evolution of I and K domains

In contrast to the previous proposition that the I domain in plant Type II (MIKC-type) TFs and the MEF2 domains in animals were independently acquired, we propose that these domains constituting the second helix in all MEF2-type TFs have a common origin. Originally, the MADS domain was defined only to include the first helix and the antiparallel strands. However, we suggest an extended definition to include the second helix, since the helix-strand-helix structure functions as one unit that probably evolved together (24, 27). Thus, the recently recognized I-like region in the Type I TFs (24) and the second helix in all other MEF2-type TFs are likely homologous, since they are structurally and functionally conserved. Meanwhile, the SAM domain in SRF-type TFs has gradually diverged from the precursor of the MEF2 domain. The turnover of the helix orientation was likely a key event establishing the two subclades, because SRF-type TFs do not heterodimerize with MEF2-type (6).

Admittedly, the predicted MEF2-type structure alone does not completely reject the previous hypothesis of an SRF-type origin of plant Type I TFs; it is possible that the hypothetically SAM-derived domain of plant Type I TFs changed the helix orientation convergently like in Type II TFs. Nevertheless, our phylogeny of the MADS domain, independent from the second helix, also suggests the MEF2-type origin of plant Type I TFs (Supplementary Fig.S3,4), which is congruent with the most parsimonious evolution of their structures.

In terms of the Keratin-like domain, we confirmed the proposed streptophytic origin of the MIKC-type (14, 15), since K domains have only been identified in streptophytic MADS-box TFs. Hence, the MEF2-type TF in the MRCA of Archaeplastida had no K domain and is thus an M-type. An ancestral M-type TF in the MRCA of Streptophyta acquired the K domain and continued evolving as a plant-specific MIKC-type. Subsequently, in the MRCA of land plants, a gene encoding MIKC-type TF duplicated into the paralogous precursors of Type I and II genes. The Type I TF precursor lost the K domain, leading to the extant Type I TFs as derived M-type.

The loss of K domain in the Type I TFs partially explains the exon number differences between Type I and II genes (8, 26). Type II genes, and most of Charophytic MADS-box genes, usually have multiple exons, with the first one coding for the MADS domain, the second one coding for the I domain and the next three to six exons coding for the K domain (8, 18, 28). Type I genes, in contrast, typically have only one or two exons, where the MADS and I-like domains are encoded by a single exon (8). The lack of introns between the MADS domain and I-like domain coding regions suggests that the Type I precursor formed through spliced mRNA and cDNA intermediates by retroposition, as previously proposed (26).

### From stress response and reproductive induction to complex structural programming

Our study also confirmed the sharp contrast between MADS-box gene family size in land plants compared to that in other eukaryotes (3, 15). Most eukaryotes have only a few MADS-box genes (Supplementary table 1), revealing that the low-copy status remained constant during the evolution of protist-like stages, including early Archaeplastida. However, following the inferred duplication of Type I and II MADS-box TFs coupled with the terrestrialization of plants, the MADS-box gene family largely expanded, which provided the raw genetic material for subsequent functional differentiation. There have been extensive studies showing that MADS-box genes are key regulators of plant organ formation, similar to homeobox genes in animals (20). The expansion of the MADS-box gene family has been proposed to be linked to the increasing complexity of extant land plants (15, 29). Convergently and in concert with the evolution of multicellularity, while less abundant in copy number, SRF and MEF2 genes in metazoan animals are both functioning in embryo patterning and continue to regulate muscle development after maturity (30). Nevertheless, since multicellularity evolved independently in animals and plants, the missing link for inferring the ancestral functional role of MADS-box genes and understanding their functional evolution lies in the unicellular, or under-differentiated multicellular eukaryotes.

Both SRF- and MEF2-type genes in unicellular and multicellular fungi, amoebae, and oomycetes have been shown to function in various stress responses (1, 31, 32, 33, 34, 35). Thus, the regulation of stress responsive programs is possibly the ancestral function of MADS-box genes, which has been maintained in multicellular metazoans, both invertebrates and vertebrates (30, 36, 37, 38). The stress-responsive rather than housekeeping function of ancestral MADS-box genes could explain the observed gene loss in several extant lineages. Originated from an ancestral stress-responsive TF, SRF- and MEF2-type TFs initially had presumably redundant functions upon duplication. Thus, the loss of SRF-type could have been compensated by MEF2-type TFs, which is likely the case in the unicellular ancestor of Archaeplastida. Supporting this assumption, the only MADS-box TF studied in microalgae, *Coccomyxa subellipsoidea* CsubMADS1, acts as a key regulator of stress tolerance (39). The colonization of the terrestrial habitat most likely required an expansion of the genetic regulators responsive to the environment. Consistently, many MADS-box TFs have known function in regulating the response to stress, like FLOWERING LOCUS C (FLC), ARABIDOPSIS NITRATE REGULATED 1 (ANR1, AGL44) or AGL21 (40).

In many unicellular organisms, the onset of reproduction is frequently induced by environmental stress, which may have facilitated the recruitment of MADS-box genes into the reproductive program (34, 35, 41, 42). In the land plant lineage, the evolution of spores and seeds that allow to withstand adverse environmental conditions may have been made possible by coupling the MADS-box TF regulation of stress resistance to reproductive development. This is particularly evident in flowering plants, where MADS-box genes regulate floral patterning, but also the onset of flowering in responses to environmental cues (40).

MADS-box TFs supposedly evolve functionally by rewiring gene regulatory networks. The DNA-binding sites recognized by SRF- and MEF2-type TFs have been extensively characterized as variants of CArG-box (6, 43), which are also conserved among investigated plant MADS-box TFs (44). Arabidopsis PHERES1 is currently the only plant Type I TF with known genome-wide DNA-binding sites *in vivo*, and the binding motifs are the same as that of Type II (45). Without dramatic sequence innovation in the DNA-binding sites, however, a given MADS-box TF achieves various collections of targets by different dimerization options (24, 46). The combination of heterodimers got largely magnified in land plants, and specifically Type I TFs no longer homodimerize (47). Moreover, the varying sequences in the C-terminus of Type I and II TFs further increased the diversity of protein-protein interaction and thus the potential to form regulatory complexes. Multiple rounds of duplication and diversification of Type I and II TFs likely have promoted the transition from a gametophyte-dominant to a sporophyte-dominant life cycle by equipping the sporophytic phase with developmental innovations such as flowers, fruits and seeds.

Interestingly, while animal MEF2 subfamily did not expand as dramatically as the land plant orthologs, large numbers of splice variants have also increased the diversity of MEF2 TFs (3, 48). Animal MEF2 TFs are expressed predominantly within the early mesoderm (30), which further differentiates into muscles, vascular and neuronal tissues, so that its function greatly promotes the mobility, integrity and sensibility of metazoans. Convergently, the plant MEF2 genes got recruited into body patterning and reproduction in response to environmental stimuli. Thus, during the evolution of multicellularity in both animals and plants, MEF2-type TFs contributed to the formation of increasingly complex body plans.

In summary, we conclude that the duplication of an ancestral MEF2-type TF with a similar domain architecture to the extant MIKC-type TFs in the MRCA of land plants gave rise to the Type I and II precursors that later progressively expanded to be an enormous family that facilitated the evolution of complex structures adapted to the terrestrial environment.

## Methods

### Sequence and phylogenetic analyses

To search for MADS-box proteins in the investigated species (Supplementary table 1), amino acid sequences of MADS-box proteins of Arabidopsis, human and yeast were used as queries in BLASTP program runs. The output sequences were aligned to the MADS domain entries in the Conserved Domain Database (49) by the conserved domain search tool, CD-Search (50), which guided the extraction of MADS domains in each species. To inspect any missed MADS-box genes, HMM searches were carried out with HMMER (51). Genomes of interest were scanned against the MADS-box TF associated profile hidden Markov model (PF00319) retrieved from Pfam (now hosted by InterPro, http://www.ebi.ac.uk/interpro/) (25).

MUSCLE was used to generate the amino acid alignments of MADS domains extracted from the selected sequences with default settings (52). We first analyzed all MADS-box TFs from a series of representative land plants: Arabidopsis (angiosperm); *Cycas panzhihuaensis*, *Ginkgo biloba* and *Gnetum luofuense* (gymnosperms); *Ceratopteris richardii* (fern), *Physcomitrium patens*, *Marchantia polymorpha* and *Anthoceros angustus* (bryophytes) (Supplementary Fig.S2). This enormous dataset was dominated by recently duplicated and possibly redundant plant sequences. Therefore, we specifically generated a downsized dataset for which the sample sizes from animals and plants were comparable, and the selected sequences of plant Type I and II TFs represented all major lineages of land plants and different well-established subfamilies. We prepared two sets of alignments for the subsequent phylogenetic analyses: the alignments of only the conventional MADS-box region corresponding the first helix and the antiparallel strands; the alignments of the extended MADS domain definition to include the second helix additionally. We also applied two different alignment tools, T-coffee (53) and MAFFT (54), which returned similar alignments (Supplementary Fig.S7).

IQ-TREE 1.6.7 was applied to perform phylogenetic analyses for maximum likelihood trees (55). The implemented ModelFinder determined LG amino acid replacement matrix (56) to be the best substitution model in the tree inference (57). 1000 replicates of ultrafast bootstraps were applied to estimate the support for reconstructed branches (58). We also selected the second-best suggestions of substitution models, JTT (59) and WAG (60), in additional ML analyses (Supplementary Fig.S8). Bayesian inference was carried out by Phylobayes (v3.2) under the CAT+GTR model with two chains. A consensus tree was built after the two chains were converged with the maxdiff less than 0.3 and the effective sample sizes of different parameters larger than 100 (61). MEGA11 (62) was applied to generate neighbour-joining trees with p-distance (proportion of different amino acids), gamma distribution allowed for rate among sites and gaps treated by pairwise deletion. 1000 bootstrap replicates were generated and majority rule defines the consensus tree.

All these alternative approaches generated phylogenetic trees in agreement with each other, reflecting the robustness of our new hypothesis. We further compared the topology of constraint phylogenetic trees fitting the previous and the new hypotheses, by several tree topology tests such as AU tests (21) supported in IQ-TREE (55).

### Protein structure prediction and analyses

We predicted the protein structures of selected MADS-box TFs by the web-based service ColabFold (https://colab.research.google.com/github/sokrypton/ColabFold/blob/main/AlphaFold2.ipynb) (63). The top ranked models were compared to resolved MADS-box protein structures, HsMEF2A (1EGW), HsSRF (1HBX) and AtSEP3 (7NB0), chains A respectively, downloaded from RCSB Protein Data Bank (https://www.rcsb.org/). The program “maxcluster” (http://www.sbg.bio.ic.ac.uk/~maxcluster/index.html) was used to perform structural comparisons based on computed TM-scores (64). Since AlphaFold2 prediction relies on MSAs, we left out nearly identical sequences which may result in the same MSAs and produce pseudo-replication. We also applied OmegaFold (https://github.com/HeliXonProtein/OmegaFold) (23) to predict the structures of all investigated MADS-box TFs based solely on their primary sequences.

## Acknowledgements

We thank Dr. Elisabeth Hehenberger for her advice on the current taxonomy of eukaryotes. This research was funded by a grant from the Knut and Alice Wallenberg Foundation (2018-0206) to C.K. and the Max Planck Society.

## Supplementary materials

**Fig.S1 a.** Bayesian inference of phylogeny of surveyed MADS domains. Posterior probabilities are labelled next to branches of interest. **b.** Neighbor-joining tree of surveyed MADS domains. Bootstrap values are labelled next to branches of interest.

**Fig.S2** Maximum likelihood tree including all MADS-box TFs from a series of representative land plants across all major lineages: Arabidopsis, *Cycas panzhihuaensis*, *Ginkgo biloba*, *Gnetum luofuense*, *Ceratopteris richardii*, *Physcomitrium patens*, *Marchantia polymorpha* and *Anthoceros angustus*. Extended MADS domain sequences were aligned by MUSCLE. The substitution model is LG. Bootstrap values are labelled next to branches of interest.

**Fig.S3** Phylogenetic trees inferred with the alignment of the conventional definition of MADS domain (only the first helix and the beta strands). **a.** Maximum likelihood tree. Bootstrap values are labelled next to branches of interest. **b.** Bayesian inference of phylogeny. Posterior probabilities are labelled next to branches of interest. **c.** Neighbor-joining tree. Bootstrap values are labelled next to branches of interest.

**Fig.S4** Topology tests for phylogenetic trees constraint to certain evolutionary models. The red tree topology is significantly better than the others. **a.** Alignments in the tests are from the same as the selected MADS-box TFs in Fig.3. Bipartition within each clade reflects the same inference as in Fig.3. **b.** Alignments in the tests are from the same as the selected MADS-box TFs in Fig.3. Internal nodes within each clade are not specified. **c.** Alignments in the tests are from the large dataset with all MADS-box TFs in representative land plants in Fig.4. Bipartition within each clade reflects the same inference as in Fig.4. **d.** Alignments in the tests are from the large dataset with all MADS-box TFs in representative land plants in Fig.4. Internal nodes within each clade are not specified.

**Fig.S5** Analyses of predicted protein structures of MADS-box transcription factors. **a-d.** Similarity scores of predicted structures in each taxonomic group to human SRF and MEF2, and Arabidopsis SEP3. The left panels are predicted by AlphaFold2, and the right panels are predicted by OmegaFold. **e.** Predicted structures of plant Type I and II proteins as monomers and dimers by AlphaFold2. Monomers are colored by confidence, red marking high confidence and blue marking low confidence. Dimers are colored to differentiate the two chains. The gray frames locate the helix-strand-helix functional units, the extended definition of MADS domains.

**Fig.S6** Maximum-Likelihood tree of surveyed MADS domains with SRF-like sequences in the core Chlorophytes (green clade pointed by the arrow). Bootstrap values are labelled above branches of interest.

**Fig.S7** Maximum likelihood trees inferred with the alignments generated by two additional softwares, T-Coffee and MAFFT. Extended MADS domain sequences were aligned. The substitution model is LG. Bootstrap values are labelled next to branches of interest. **a.** T-coffee. **b.** MAFFT.

**Fig.S8** Maximum likelihood trees inferred with two additionally substitution models, JTT and WAG. The alignments of extended MADS domain sequences were generated by MUSCLE. Bootstrap values are labelled next to branches of interest. **a.** JTT. **b.** WAG.

**Table S1.** Species surveyed in this study.

**Table S2.** Protein similarity scores of AlphaFold2 predicted models for MADS-box TFs to human SRF and MEF2, and Arabidopsis AtSEP3.

**Table S3.** Protein similarity scores of OmegaFold predicted models for MADS-box TFs to human SRF and MEF2, and Arabidopsis AtSEP3.

## References

1. F. Messenguy, E. Dubois (2003) Role of MADS box proteins and their cofactors in combinatorial control of gene expression and cell development. Gene 316, 1–21.

2. Z. Schwarz-Sommer, P. Huijser, W. Nacken, H. Saedler, H. Sommer (1990) Genetic control of flower development by homeotic genes in Antirrhinum majus. Science 250, 931–936.

3. G. Theissen, J. T. Kim, H. Saedler (1996) Classification and phylogeny of the MADS-box multigene family suggest defined roles of MADS-box gene subfamilies in the morphological evolution of eukaryotes. J. Mol. Evol. 43, 484–516.

4. E. R. Alvarez-Buylla et al. (2000) An ancestral MADS-box gene duplication occurred before the divergence of plants and animals. Proc. Natl. Acad. Sci. U.S.A. 97, 5328–5333.

5. L. Gramzow, M. S. Ritz, G. Theissen (2010) On the origin of MADS-domain transcription factors. Trends Genet. 26, 149–153.

6. P. Shore, A. D. Sharrocks (1995) The MADS-box family of transcription factors. Eur. J. Biochem. 229, 1–13.

7. L. Gramzow, G. Theissen (2013) Phylogenomics of MADS-Box genes in plants-two opposing life styles in one gene family. Biology (Basel) 2, 1150–1164.

8. L. Parenicová et al. (2003) Molecular and phylogenetic analyses of the complete MADS-box transcription factor family in Arabidopsis: new openings to the MADS world. Plant Cell 15, 1538–1551.

9. M. Bemer, K. Heijmans, C. Airoldi, B. Davies, G. C. Angenent (2010) An atlas of type I MADS box gene expression during female gametophyte and seed development in Arabidopsis. Plant Physiol. 154, 287–300.

10. S. Masiero, L. Colombo, P. E. Grini, A. Schnittger, M. M. Kater (2011) The emerging importance of type I MADS box transcription factors for plant reproduction. Plant Cell 23, 865–872.

11. Y. Qiu, C. Köhler (2022) Endosperm evolution by duplicated and neofunctionalized Type I MADS-box transcription factors. Mol. Biol. Evol. 39, msab355.

12. J. Nam et al. (2004) Type I MADS-box genes have experienced faster birth-and-death evolution than type II MADS-box genes in angiosperms. Proc. Natl. Acad. Sci. U.S.A. 101, 1910–1915.

13. M. Ng, M. F. Yanofsky (2001). Function and evolution of the plant MADS-box gene family. Nat. Rev. Genet. 2, 186–195.

14. K. Kaufmann, R. Melzer, G. Theissen (2005) MIKC-type MADS-domain proteins: structural modularity, protein interactions and network evolution in land plants. Gene 347, 183–198.

15. G. Thangavel, and S. Nayar (2018) A survey of MIKC Type MADS-box genes in non-seed plants: Algae, Bryophytes, Lycophytes and Ferns. Front. Plant Sci. 9, 510.

16. S. De Bodt, J. Raes, Y. Van de Peer, G. Theissen (2003) And then there were many: MADS goes genomic. Trends Plant Sci. 8, 475–483.

17. K. Hori et al. (2014) Klebsormidium flaccidum genome reveals primary factors for plant terrestrial adaptation. Nat. Commun. 5, 3978.

18. T. Nishiyama et al. (2018) The Chara genome: secondary complexity and implications for plant terrestrialization. Cell 174, 448–464.e24.

19. F. Burki, A. J. Roger, M. W. Brown, A. Simpson (2020) The new tree of eukaryotes. Trends Ecol. Evol. 35, 43–55.

20. J. Nam, C. W. dePamphilis, H. Ma, M. Nei (2003) Antiquity and evolution of the MADS-box gene family controlling flower development in plants. Mol. Biol. Evol. 20, 1435–1447.

21. H. Shimodaira (2002) An approximately unbiased test of phylogenetic tree selection. Syst. Biol. 51, 492–508.

22. J. Jumper et al. (2021) Highly accurate protein structure prediction with AlphaFold. Nature 596 583–589.

23. R. Wu et al. (2022) High-resolution de novo structure prediction from primary sequence. bioRxiv https://www.biorxiv.org/content/10.1101/2022.07.21.500999v1.

24. X. Lai et al. (2021) The intervening domain is required for DNA-binding and functional identity of plant MADS transcription factors. Nat. Commun. 12, 4760.

25. T. Paysan-Lafosse et al. (2023) InterPro in 2022. Nucleic Acids Res. 51(D1), D418– D427.

26. R. Kofuji et al. (2003) Evolution and divergence of the MADS-box gene family based on genome-wide expression analyses. Mol. Biol. Evol. 20, 1963–1977.

27. J. L. Riechmann, B. A. Krizek, E. M. Meyerowitz (1996) Dimerization specificity of Arabidopsis MADS domain homeotic proteins APETALA1, APETALA3, PISTILLATA, and AGAMOUS. Proc. Natl. Acad. Sci. U.S.A. 93, 4793–4798.

28. F. Rümpler et al. (2023) The Origin of Floral Quartet Formation-Ancient Exon Duplications Shaped the Evolution of MIKC-type MADS-domain Transcription Factor Interactions. Mol. Biol. Evol. 40, msad088.

29. L. Gramzow, L. Weilandt, G. Theissen (2014) MADS goes genomic in conifers: towards determining the ancestral set of MADS-box genes in seed plants. Ann. Bot. 114, 1407–1429.

30. M. Potthoff, E. N. Olson (2007) MEF2: a central regulator of diverse developmental programs. Development 134, 4131–4140.

31. Z. Ding, T. Xu, W. Zhu, L. Li, Q. Fu (2020) A MADS-box transcription factor FoRlm1 regulates aerial hyphal growth, oxidative stress, cell wall biosynthesis and virulence in Fusarium oxysporum f. sp. cubense. Fungal Biol. 124, 183–193.

32. M. C. Rocha et al. (2016) Aspergillus fumigatus MADS-box transcription factor rlmA is required for regulation of the cell wall integrity and virulence. G3 (Bethesda) 6, 2983–3002.

33. Q. Wang et al. (2018) MADS-box transcription factor MadsA regulates dimorphic transition, conidiation, and germination of Talaromyces marneffei. Front. Microbiol. 9, 1781.

34. M. Galardi-Castilla, I. Fernandez-Aguado, T. Suarez, L. Sastre (2013) Mef2A, a homologue of animal Mef2 transcription factors, regulates cell differentiation in Dictyostelium discoideum. BMC Dev. Biol. 13, 12.

35. W. Leesutthiphonchai, H. S. Judelson (2018) A MADS-box transcription factor regulates a central step in sporulation of the oomycete Phytophthora infestans. Mol. Microbiol. 110, 562–575.

36. A. Vrailas-Mortimer et al. (2011) A muscle-specific p38 MAPK/Mef2/MnSOD pathway regulates stress, motor function, and life span in Drosophila. Dev. Cell 21, 783–795.

37. F. J. Blanchard et al. (2010) The transcription factor Mef2 is required for normal circadian behavior in Drosophila. J. Neurosci. 30, 5855–5865.

38. A. M. van der Linden, K. M. Nolan, P. Sengupta (2007) KIN-29 SIK regulates chemoreceptor gene expression via an MEF2 transcription factor and a class II HDAC. EMBO J. 26, 358–370.

39. S. Nayar, G. Thangavel (2021) CsubMADS1, a lag phase transcription factor, controls development of polar eukaryotic microalga Coccomyxa subellipsoidea C-169. Plant J. 107, 1228–1242.

40. N. Castelán-Muñoz et al. (2019) MADS-box genes are key components of genetic regulatory networks involved in abiotic stress and plastic developmental responses in plants. Front. Plant Sci. 10, 853.

41. S. Piccirillo et al. (2015) Cell differentiation and spatial organization in yeast colonies: role of cell-wall integrity pathway. Genetics 201, 1427–1438.

42. R. Escalante, N. Moreno, L. Sastre (2003) Dictyostelium discoideum developmentally regulated genes whose expression is dependent on MADS box transcription factor SrfA. Eukaryot. Cell 2, 1327–1335.

43. W. Wu et al. (2011) Conservation and evolution in and among SRF- and MEF2- type MADS domains and their binding sites. Mol. Biol. Evol. 28, 501–511.

44. N. Aerts, S. de Bruijn, H. van Mourik, G. C. Angenent, A. D. J. van Dijk (2018) Comparative analysis of binding patterns of MADS-domain proteins in Arabidopsis thaliana. BMC Plant Biol. 18, 131.

45. R. A. Batista et al. (2019) The MADS-box transcription factor PHERES1 controls imprinting in the endosperm by binding to domesticated transposons. eLife 8, e50541.

46. H. van Mourik et al. (2023) Dual specificity and target gene selection by the MADS-domain protein FRUITFULL. Nat. Plants 9, 473–485.

47. S. de Folter et al. (2005) Comprehensive interaction map of the Arabidopsis MADS Box transcription factors. Plant Cell 17, 1424–1433.

48. J. F. Martin et al. (1994) A Mef2 gene that generates a muscle-specific isoform via alternative mRNA splicing. Mol. Cell Biol. 14, 1647–1656.

49. S. Lu, et al. (2020) CDD/SPARCLE: the conserved domain database in 2020. Nucleic Acids Res. 48(D1), D265-D268.

50. A. Marchler-Bauer, S. H. Bryant (2004) CD-Search: protein domain annotations on the fly. Nucleic Acids Res. 32, W327–W331.

51. S. R. Eddy (2011) Accelerated profile HMM searches. PLoS Comput. Biol. 7, e1002195.

52. R. C. Edgar (2004) MUSCLE: multiple sequence alignment with high accuracy and high throughput. Nucleic Acids Res. 32, 1792–1797.

53. C. Notredame, D. G. Higgins, J. Heringa (2000) T-Coffee: A novel method for fast and accurate multiple sequence alignment. J. Mol. Biol. 302, 205–217.

54. K. Katoh, J. Rozewicki, K. D. Yamada (2019) MAFFT online service: multiple sequence alignment, interactive sequence choice and visualization. Brief. Bioinform. 20, 1160–1166.

55. L. T. Nguyen, H. A. Schmidt, A. von Haeseler, B. Q. Minh (2015) IQ-TREE: a fast and effective stochastic algorithm for estimating maximum-likelihood phylogenies. Mol. Biol. Evol. 32, 268–274.

56. S. Q. Le, O. Gascuel (2008) An improved general amino acid replacement matrix. Mol. Biol. Evol. 25, 1307–1320.

57. S. Kalyaanamoorthy, B. Q. Minh, T. Wong, A. von Haeseler, L. S. Jermiin (2017) ModelFinder: fast model selection for accurate phylogenetic estimates. Nat. Methods 14, 587–589.

58. D. T. Hoang, O. Chernomor, A. von Haeseler, B. Q. Minh, L. S. Vinh (2018) UFBoot2: improving the ultrafast bootstrap approximation. Mol. Biol. Evol. 35, 518–522.

59. D. T. Jones, W. R. Taylor, J. M. Thornton (1992) The rapid generation of mutation data matrices from protein sequences. Bioinformatics 8, 275–282.

60. S. Whelan, N. Goldman (2001) A general empirical model of protein evolution derived from multiple protein families using a maximum-likelihood approach. Mol. Biol. Evol. 18, 691–699.

61. N. Lartillot, T. Lepage, S. Blanquart (2009) PhyloBayes 3: a Bayesian software package for phylogenetic reconstruction and molecular dating. Bioinformatics 25, 2286–2288.

62. K. Tamura, G. Stecher, S. Kumar (2021) MEGA11: Molecular Evolutionary Genetics Analysis Version 11. Mol. Biol. Evol. 38, 3022–3027.

63. M. Mirdita et al. (2022) ColabFold: making protein folding accessible to all. Nat. Methods 19, 679–682.

64. Y. Zhang, J. Skolnick (2005) TM-align: a protein structure alignment algorithm based on the TM-score. Nucleic Acids Res. 33, 2302–2309.

65. L. Schrödinger, W. DeLano (2020) PyMOL. http://www.pymol.org/pymol.

