## Supplemental Figures for "Updated phylogeny and protein structure predictions revise the hypothesis on the origin of MADS-box transcription factors in land plants"

**Fig.S1 a.** Bayesian inference of phylogeny of surveyed MADS domains. Posterior probabilities are labelled next to branches of interest.

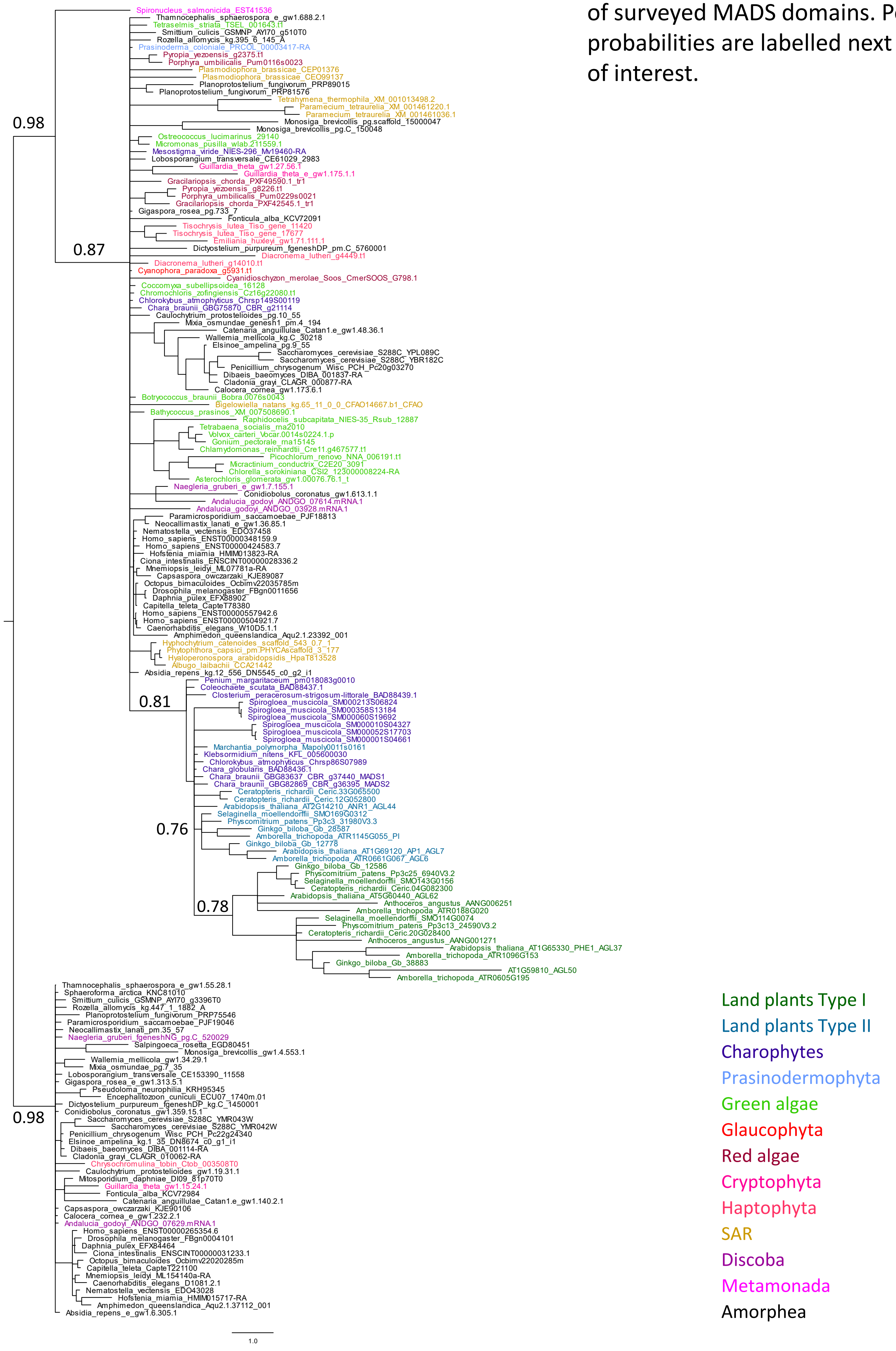

**Fig.S1 b.** Neighbor-joining tree of surveyed MADS domains. Bootstrap values are labelled next to branches of interest.

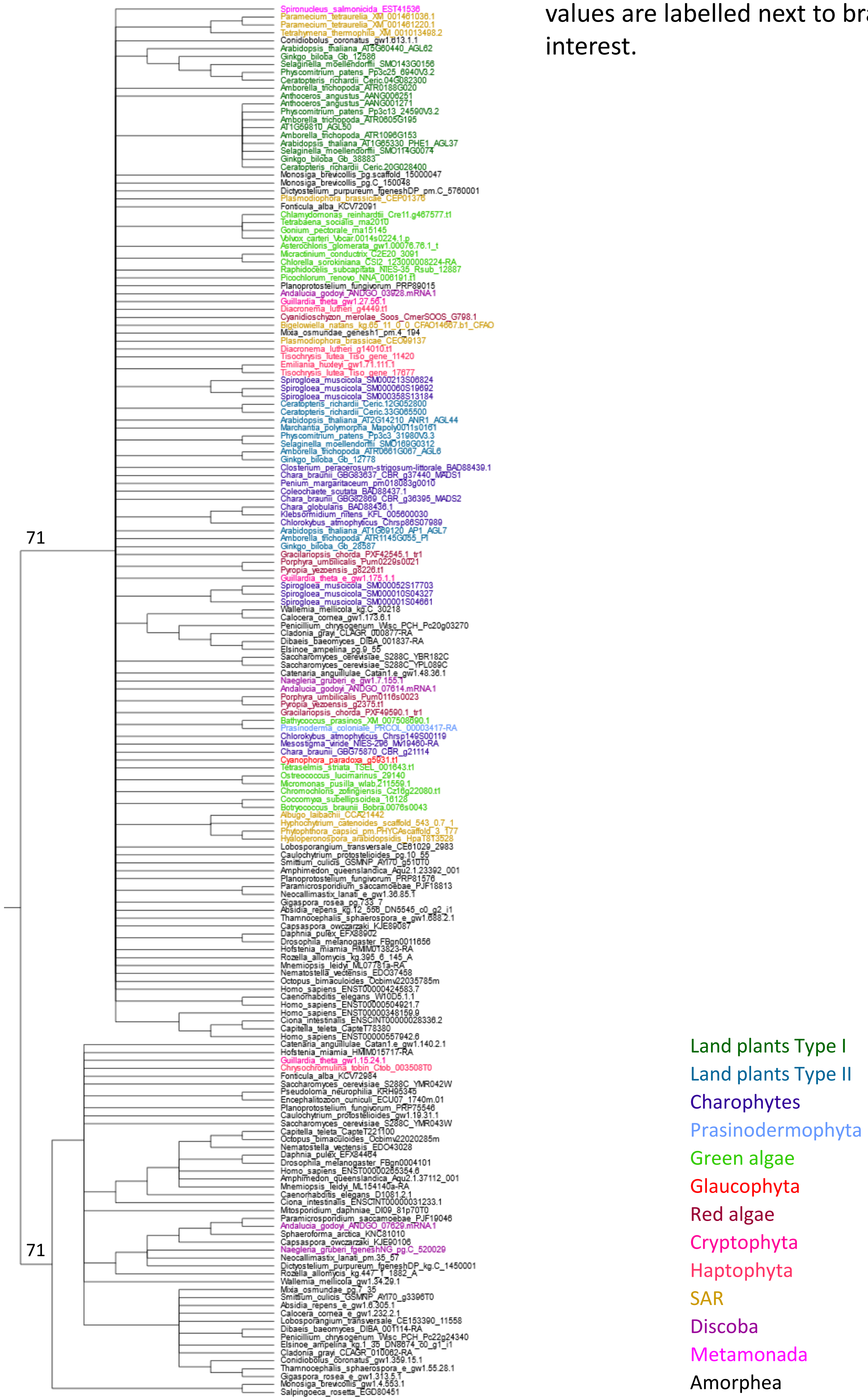

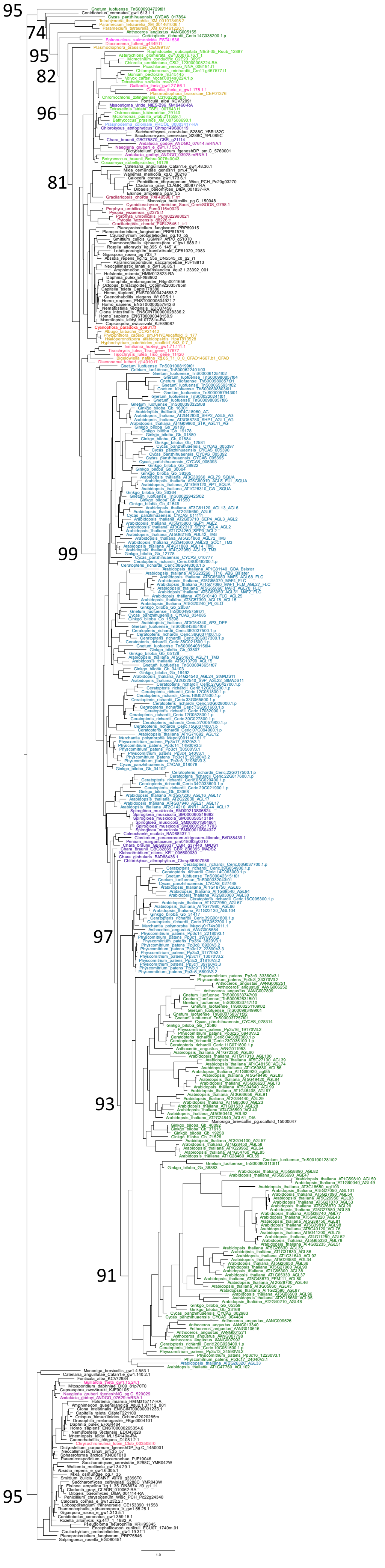


**Fig.S2** Maximum likelihood tree including all MADS-box TFs from a series of representative land plants across all major lineages: Arabidopsis, *Cycas panzhihuaensis*, *Ginkgo biloba*, *Gnetum luofuense*, *Ceratopteris richardii*, *Physcomitrium patens*, *Marchantia polymorpha* and *Anthoceros angustus*. Extended MADS domain sequences were aligned by MUSCLE. The substitution model is LG. Bootstrap values are labelled next to branches of interest.

**Fig.S3** Phylogenetic trees inferred with the alignment of the conventional definition of MADS domain (only the first helix and the beta strands).

**a.** Maximum likelihood tree. Bootstrap values are labelled next to branches of interest.

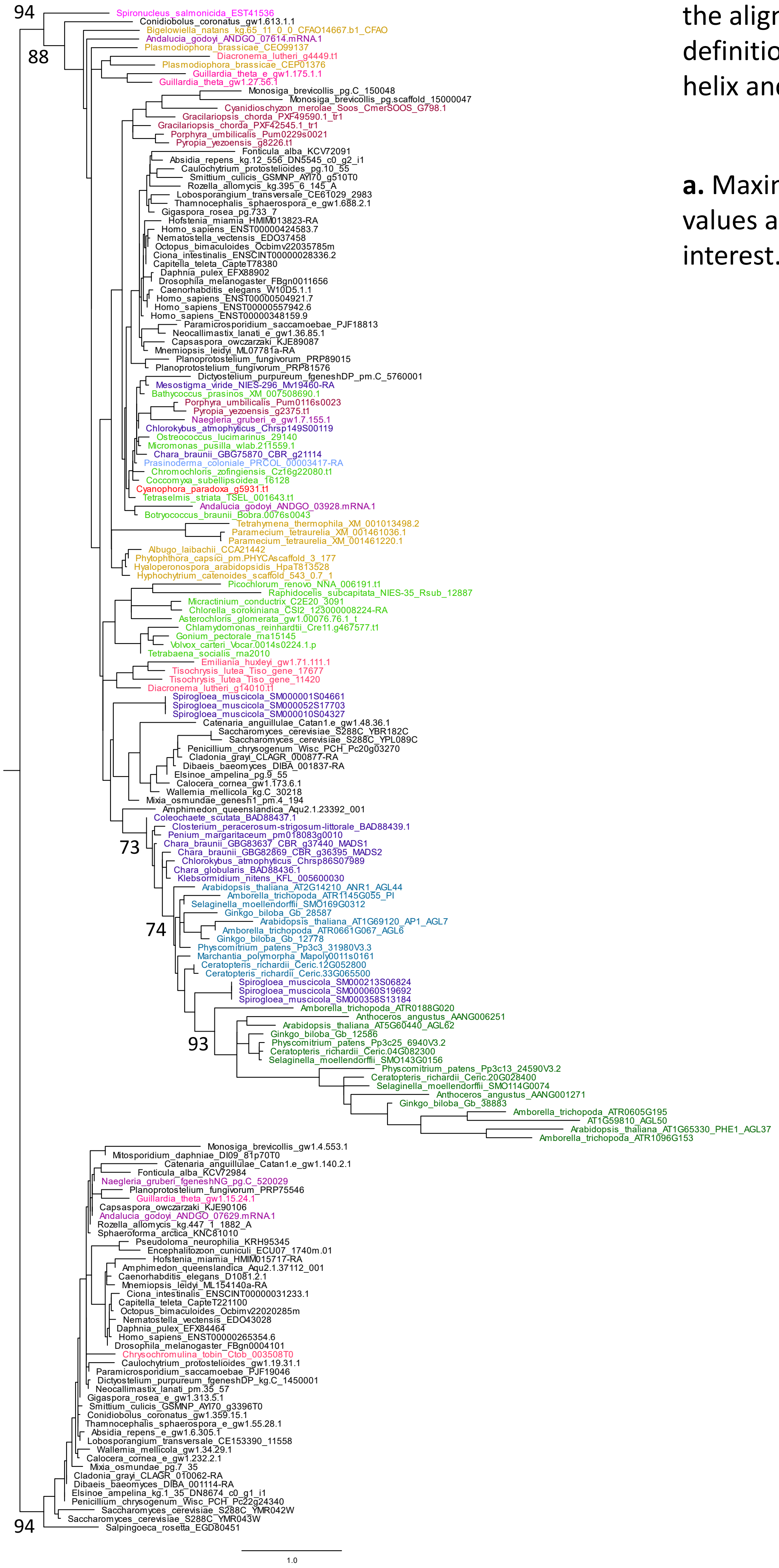

**Fig.S3** Phylogenetic trees inferred with the alignment of the conventional definition of MADS domain (only the first helix and the beta strands).

**b.** Bayesian inference of phylogeny. Posterior probabilities are labelled next to branches of interest.

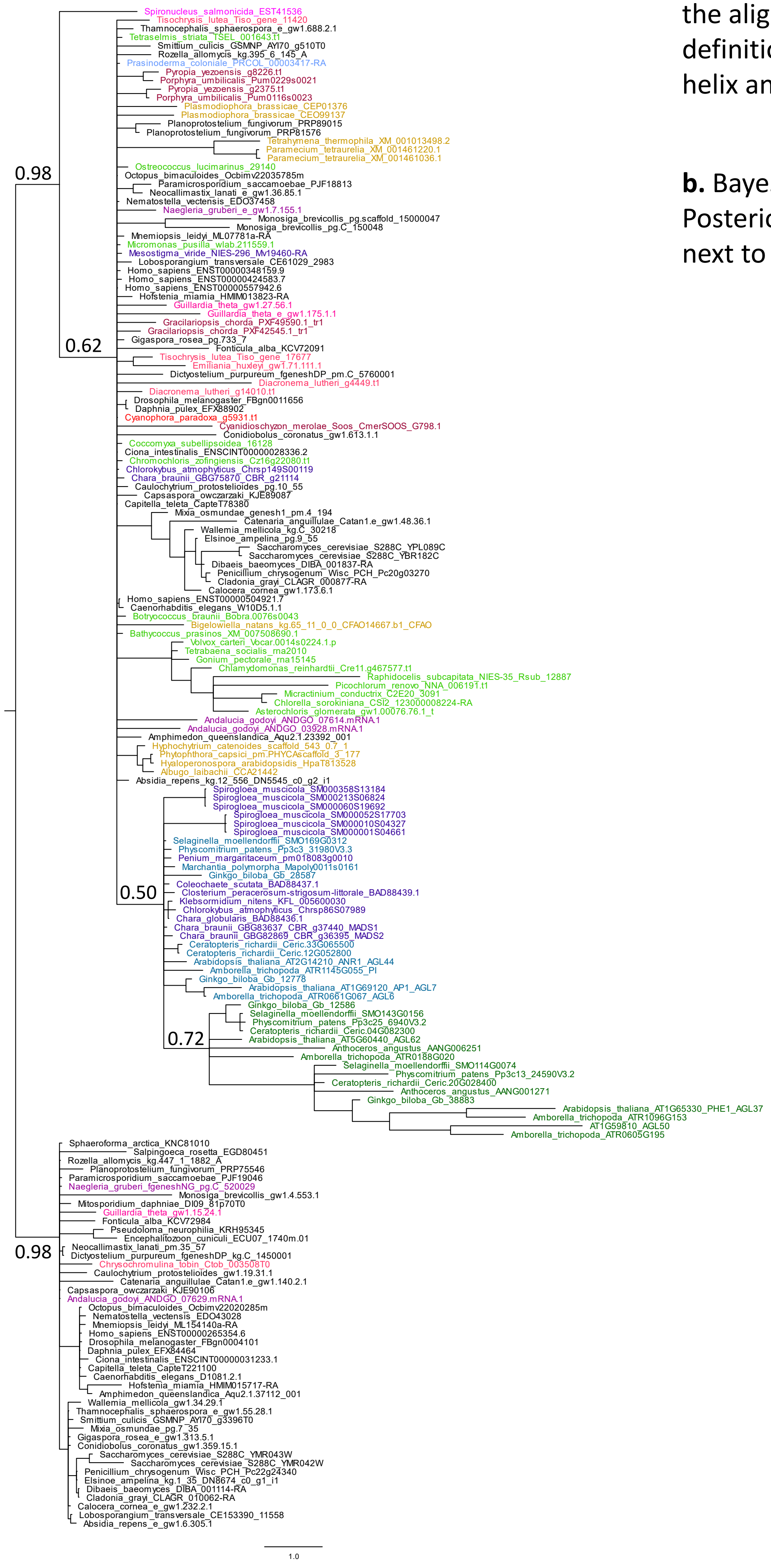

**Fig.S3** Phylogenetic trees inferred with the alignment of the conventional definition of MADS domain (only the first helix and the beta strands).

**c.** Neighbor-joining tree. Bootstrap values are labelled next to branches of interest.

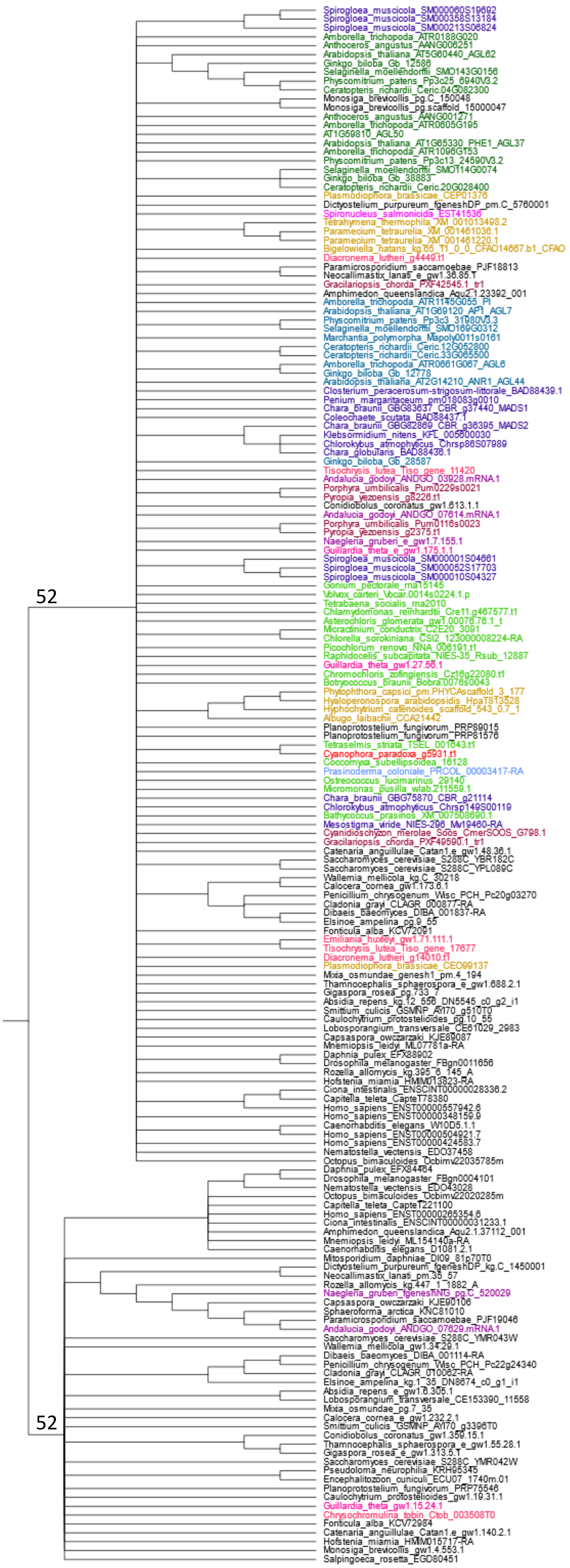


Land plants Type I
Land plants Type II
Charophytes
Prasinodermophyta
Green algae
Glaucophyta
Red algae
Cryptophyta
Haptophyta
SAR
Discoba
Metamonada
Amorphea

**Fig.S4** Topology tests for phylogenetic trees constraint to certain evolutionary models. The red tree topology is significantly better than the others.

**a.** Alignments in the tests are from the same as the selected MADS-box TFs in Fig.3. Bipartition within each clade reflects the same inference as in Fig.3.

Test 1: Input was the alignments of the extended MADS domain (helix-strands-helix)

| Tree | logL | deltaL | bp-RELL | | p-KH | | p-SH | | p-WKH | | p-WSH | | c-ELW | | p-AU | |
|---|---|---|---|---|---|---|---|---|---|---|---|---|---|---|---|---|
| 1 | -15250.99413 | 41.52 | 0.0001 | - | 0.0008 | - | 0.0008 | - | 0.0008 | - | 0.0008 | - | 0.00017 | - | 6.7e-05 | - |
| 2 | -15209.47455 | 0 | 1 | + | 0.999 | + | 1 | + | 0.999 | + | 0.999 | + | 1 | + | 1 | + |

Test 2: Input was the alignments of the conventional MADS domain (helix-strands)

| Tree | logL | deltaL | bp-RELL | | p-KH | | p-SH | | p-WKH | | p-WSH | | c-ELW | | p-AU | |
|---|---|---|---|---|---|---|---|---|---|---|---|---|---|---|---|---|
| 1 | -7757.881432 | 32.875 | 0.0008 | - | 0.0034 | - | 0.0034 | - | 0.0034 | - | 0.0034 | - | 0.00107 | - | 0.000618 | - |
| 2 | -7725.006562 | 0 | 0.999 | + | 0.997 | + | 1 | + | 0.997 | + | 0.997 | + | 0.999 | + | 0.999 | + |

Stats of tree topology tests from IQ-tree.

```
deltaL  : logL difference from the maximal logl in the set.
bp-RELL : bootstrap proportion using RELL method (Kishino et al. 1990).
p-KH    : p-value of one sided Kishino-Hasegawa test (1989).
p-SH    : p-value of Shimodaira-Hasegawa test (2000).
p-WKH   : p-value of weighted KH test.
p-WSH   : p-value of weighted SH test.
c-ELW   : Expected Likelihood Weight (Strimmer & Rambaut 2002).
p-AU    : p-value of approximately unbiased (AU) test (Shimodaira, 2002).

Plus signs denote the 95% confidence sets (possible topology).
Minus signs denote significant exclusion (rejected topology).
```

Tree 1
Hypothesis by Alvarez-Buylla et al. (2000):
Land plant Type I is SRF-type, Type II is MEF2-type.

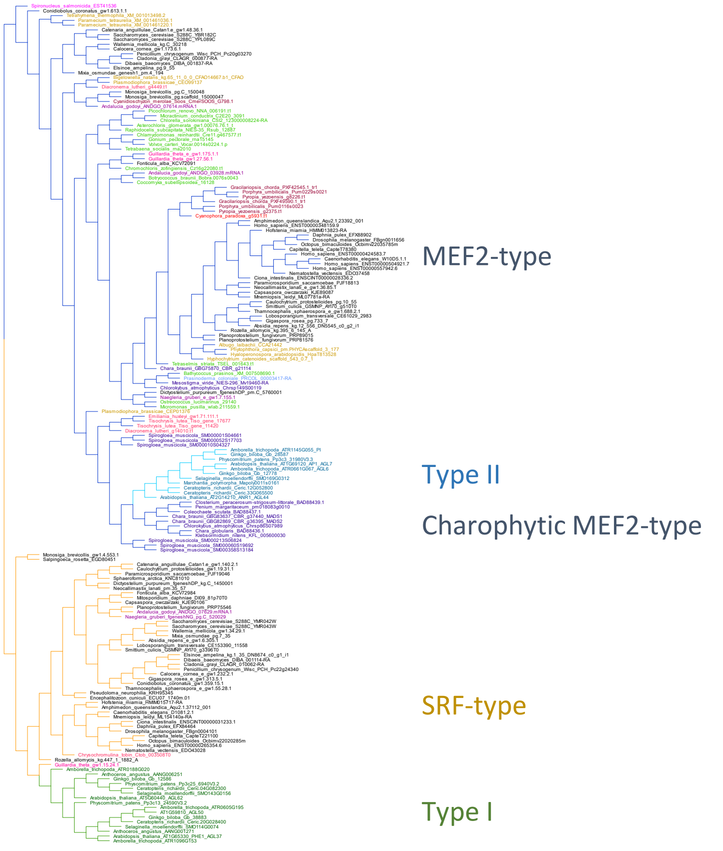


Tree 2
Hypothesis by this study:
Land plant Type I and II both are MEF2-type.

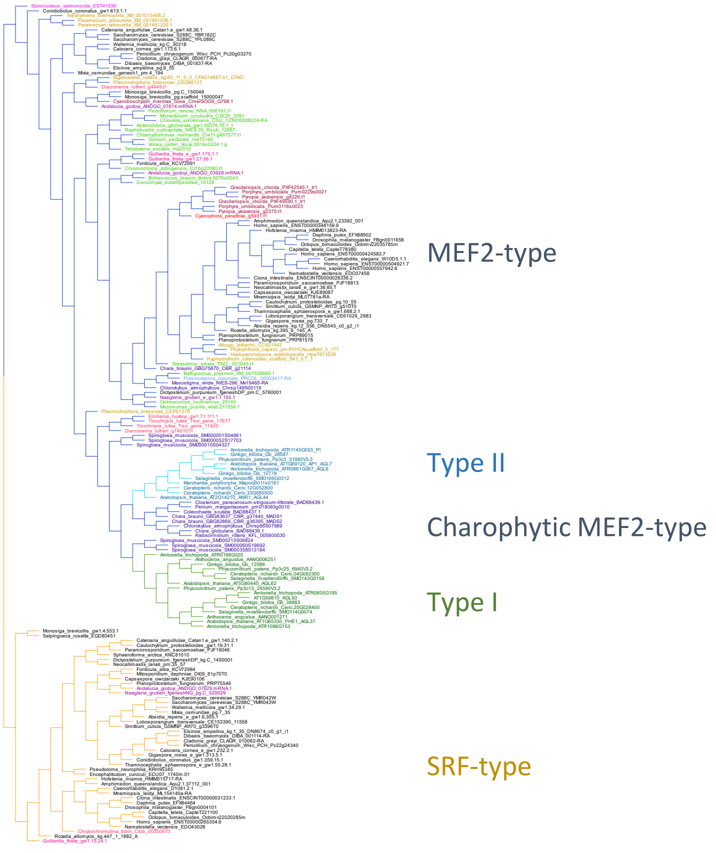

**Fig.S4** Topology tests for phylogenetic trees constraint to certain evolutionary models. The red tree topology is significantly better than the others.

**b.** Alignments in the tests are from the same as the selected MADS-box TFs in Fig.3. Internal nodes within each clade are not specified.

Test 1: Input was the alignments of the extended MADS domain (helix-strands-helix)

| Tree | logL | deltaL | bp-RELL | | p-KH | | p-SH | | p-WKH | | p-WSH | | c-ELW | | p-AU | |
|---|---|---|---|---|---|---|---|---|---|---|---|---|---|---|---|---|
| 1 | -19283.08554 | 98.415 | 0 | - | 0 | - | 0 | - | 0 | - | 0 | - | 8.27e-27 | - | 6.51e-05 | - |
| 2 | -19272.28884 | 87.619 | 0 | - | 0 | - | 0 | - | 0 | - | 0 | - | 7.26e-20 | - | 6.27e-51 | - |
| 3 | -19225.18385 | 40.514 | 0.0005 | - | 0.0011 | - | 0.0091 | - | 0.0011 | - | 0.0015 | - | 0.000483 | - | 0.000158 | - |
| 4 | -19184.67007 | 0 | 1 | + | 0.999 | + | 1 | + | 0.999 | + | 1 | + | 1 | + | 1 | + |

Test 2: Input was the alignments of the conventional MADS domain (helix-strands)

| Tree | logL | deltaL | bp-RELL | | p-KH | | p-SH | | p-WKH | | p-WSH | | c-ELW | | p-AU | |
|---|---|---|---|---|---|---|---|---|---|---|---|---|---|---|---|---|
| 1 | -10677.37393 | 65.977 | 0 | - | 0 | - | 0 | - | 0 | - | 0 | - | 8.51e-15 | - | 3.45e-08 | - |
| 2 | -10678.09232 | 66.696 | 0 | - | 0 | - | 0 | - | 0 | - | 0 | - | 2.23e-13 | - | 1.34e-49 | - |
| 3 | -10640.48744 | 29.091 | 0.0022 | - | 0.0036 | - | 0.0111 | - | 0.0036 | - | 0.0073 | - | 0.00253 | - | 0.000954 | - |
| 4 | -10611.39666 | 0 | 0.998 | + | 0.996 | + | 1 | + | 0.996 | + | 1 | + | 0.997 | + | 0.999 | + |

Tree 1
Hypothesis not proposed ever:
Land plant Type I and II both are SRF-type.

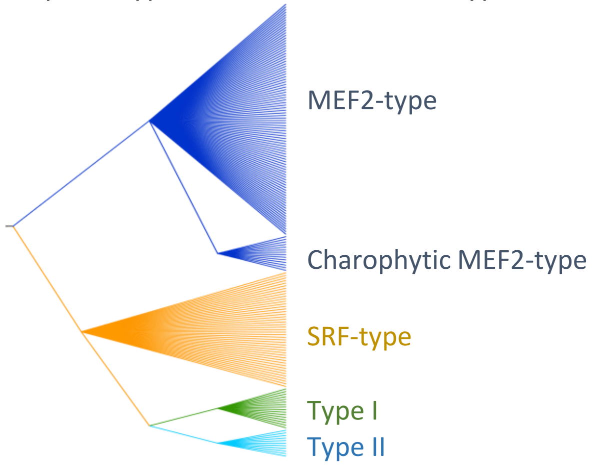


Tree 4
Hypothesis not proposed ever:
Land plant Type I is MEF2-type, Type II is SRF-type.

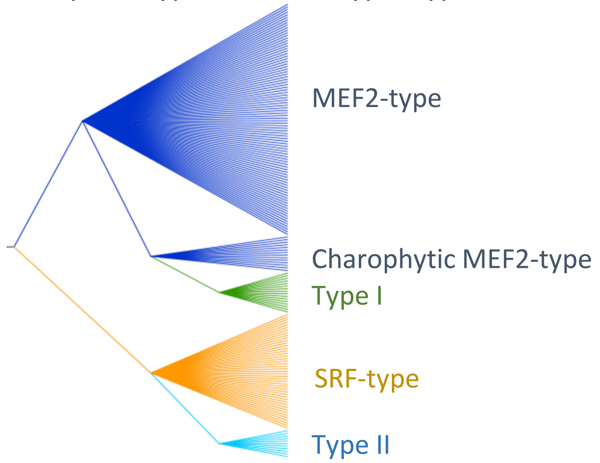


Tree 3
Hypothesis by Alvarez-Buylla et al. (2000):
Land plant Type I is SRF-type, Type II is MEF2-type.

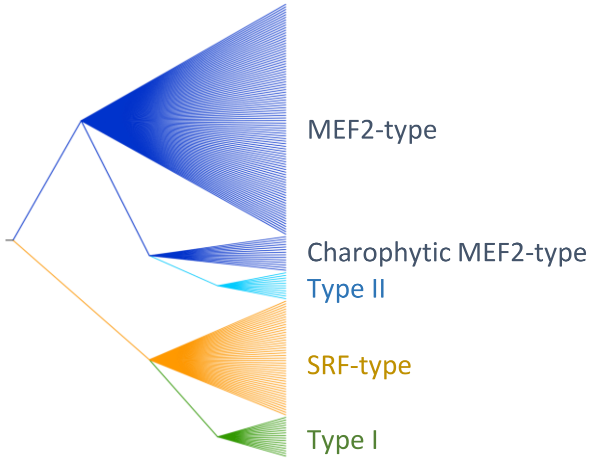


Tree 4
Hypothesis by this study:
Land plant Type I and II both are MEF2-type.

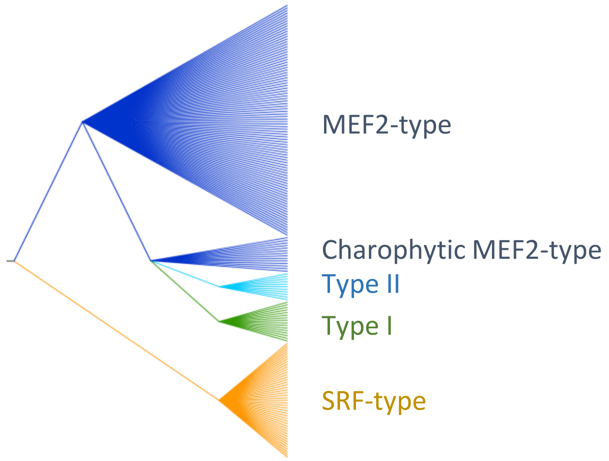

**Fig.S4** Topology tests for phylogenetic trees constraint to certain evolutionary models. The red tree topology is significantly better than the others.

**c.** Alignments in the tests are from the large dataset with all MADS-box TFs in representative land plants in Fig.4. Bipartition within each clade reflects the same inference as in Fig.4.

Input was the alignments of the extended MADS domain (helix-strands-helix)

| Tree | logL | deltaL | bp-RELL | | p-KH | | p-SH | | p-WKH | | p-WSH | | c-ELW | | p-AU | |
|---|---|---|---|---|---|---|---|---|---|---|---|---|---|---|---|---|
| 1 | -32494.46232 | 52.521 | 0.0006 | - | 0.0019 | - | 0.0019 | - | 0.0019 | - | 0.0019 | - | 0.000637 | - | 0.000353 | - |
| 2 | -32441.94179 | 0 | 0.999 | + | 0.998 | + | 1 | + | 0.998 | + | 0.998 | + | 0.999 | + | 1 | + |

Stats of tree topology tests from IQ-tree.

```
deltaL  : logL difference from the maximal logl in the set.
bp-RELL : bootstrap proportion using RELL method (Kishino et al. 1990).
p-KH    : p-value of one sided Kishino-Hasegawa test (1989).
p-SH    : p-value of Shimodaira-Hasegawa test (2000).
p-WKH   : p-value of weighted KH test.
p-WSH   : p-value of weighted SH test.
c-ELW   : Expected Likelihood Weight (Strimmer & Rambaut 2002).
p-AU    : p-value of approximately unbiased (AU) test (Shimodaira, 2002).

Plus signs denote the 95% confidence sets (possible topology).
Minus signs denote significant exclusion (rejected topology).
```

Tree 1
Hypothesis by Alvarez-Buylla et al. (2000):
Land plant Type I is SRF-type, Type II is MEF2-type.

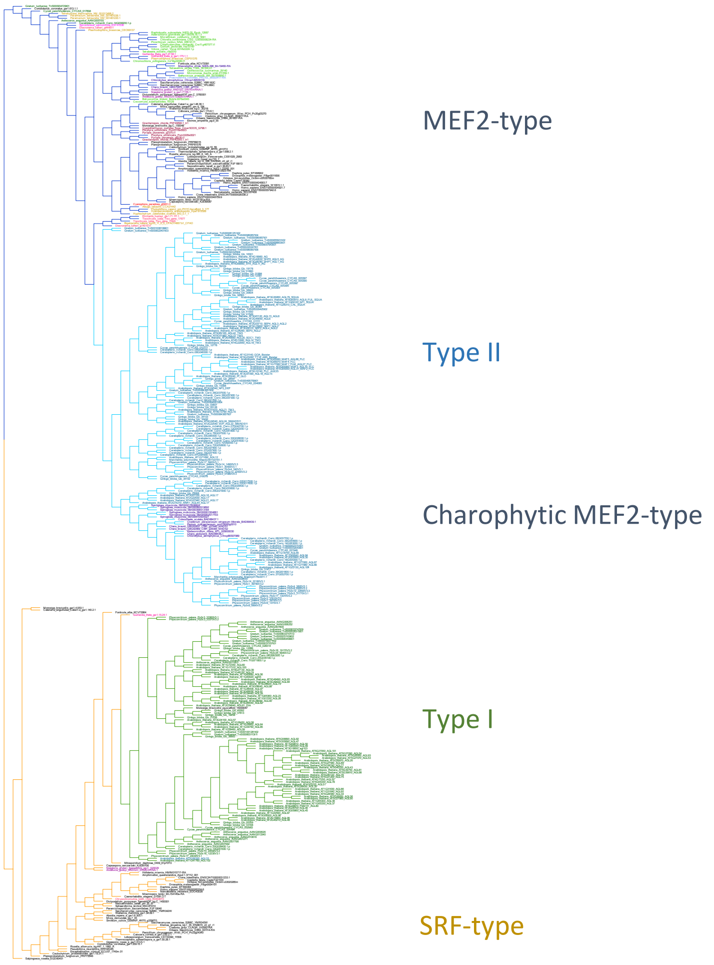


Tree 2
Hypothesis by this study:
Land plant Type I and II both are MEF2-type.

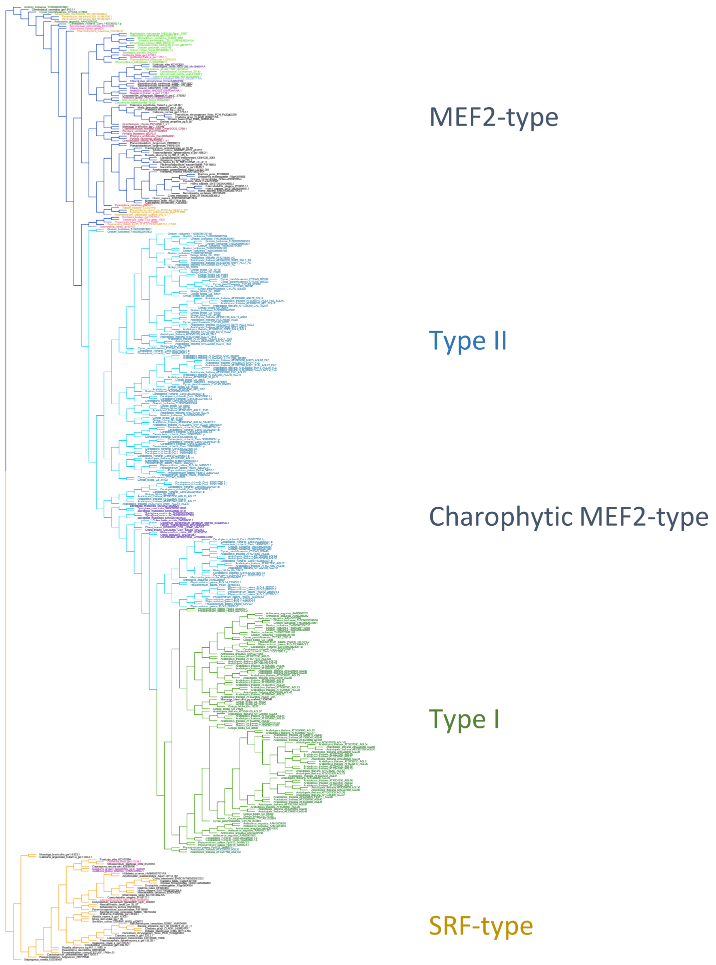

**Fig.S4** Topology tests for phylogenetic trees constraint to certain evolutionary models. The red tree topology is significantly better than the others.

**d.** Alignments in the tests are from the large dataset with all MADS-box TFs in representative land plants in Fig.4. Internal nodes within each clade are not specified.

Input was the alignments of the extended MADS domain (helix-strands-helix)

| Tree | logL | deltaL | bp-RELL | | p-KH | | p-SH | | p-WKH | | p-WSH | | c-ELW | | p-AU | |
|---|---|---|---|---|---|---|---|---|---|---|---|---|---|---|---|---|
| 1 | -48257.98946 | 33.929 | 0.0044 | - | 0.0043 | - | 0.0043 | - | 0.0043 | - | 0.0043 | - | 0.00466 | - | 0.00373 | - |
| 2 | -48224.06043 | 0 | 0.996 | + | 0.996 | + | 1 | + | 0.996 | + | 0.996 | + | 0.995 | + | 0.996 | + |

Tree 1
Hypothesis by Alvarez-Buylla et al. (2000):
Land plant Type I is SRF-type, Type II is MEF2-type.

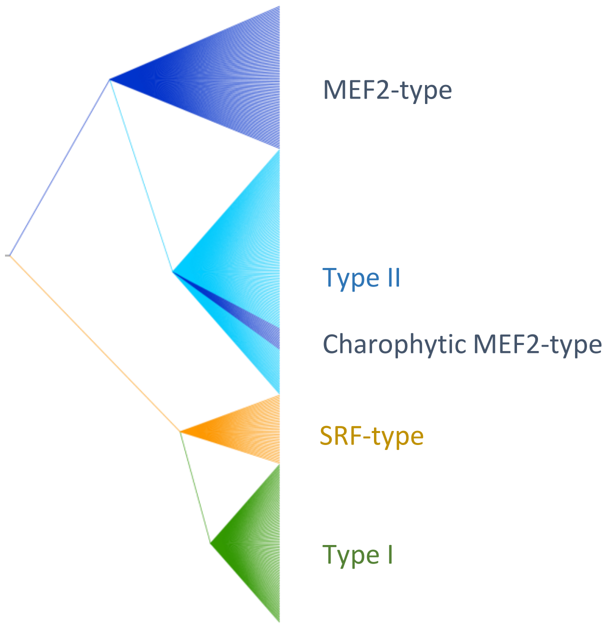


Tree 2
Hypothesis by this study:
Land plant Type I and II both are MEF2-type.

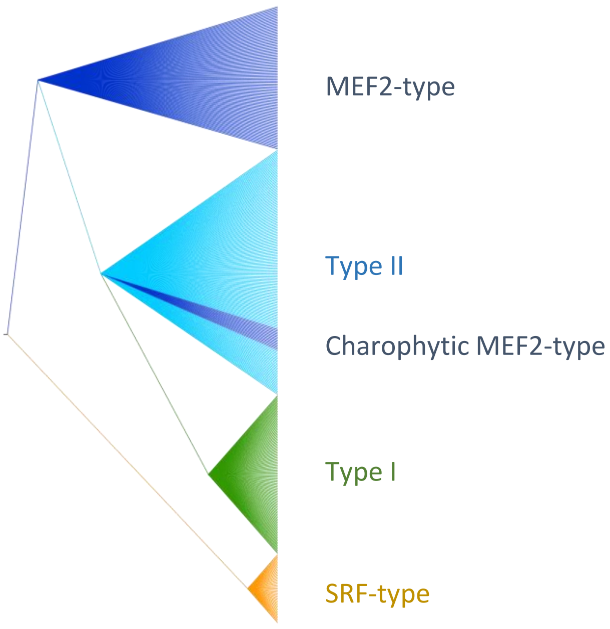

**Fig.S5** Analyses of predicted protein structures of MADS-box transcription factors.

**a-d.** Similarity scores of predicted structures in each taxonomic group to human SRF and MEF2, and Arabidopsis SEP3. The left panels are predicted by AlphaFold2, and the right panels are predicted by OmegaFold.

**a.** Amorphea, Discoba and Metamonada

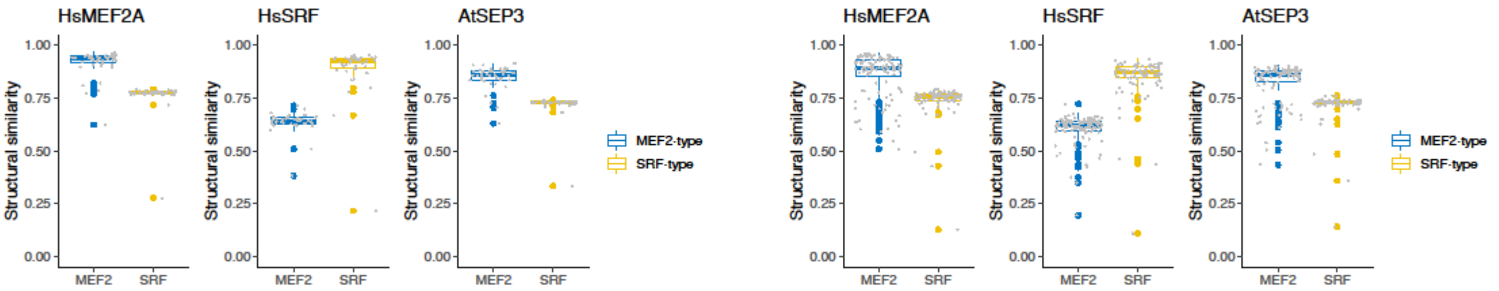


**b.** Non-streptophytic Archaeplastida

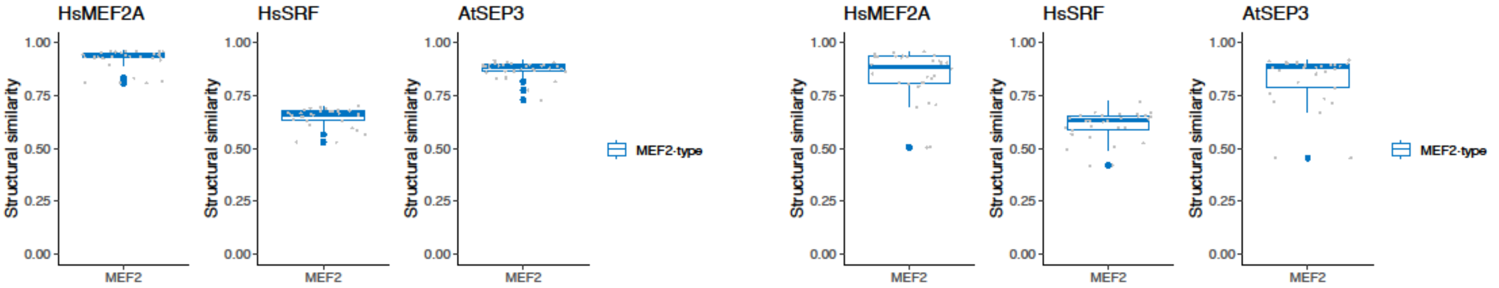


**c.** Charophytes

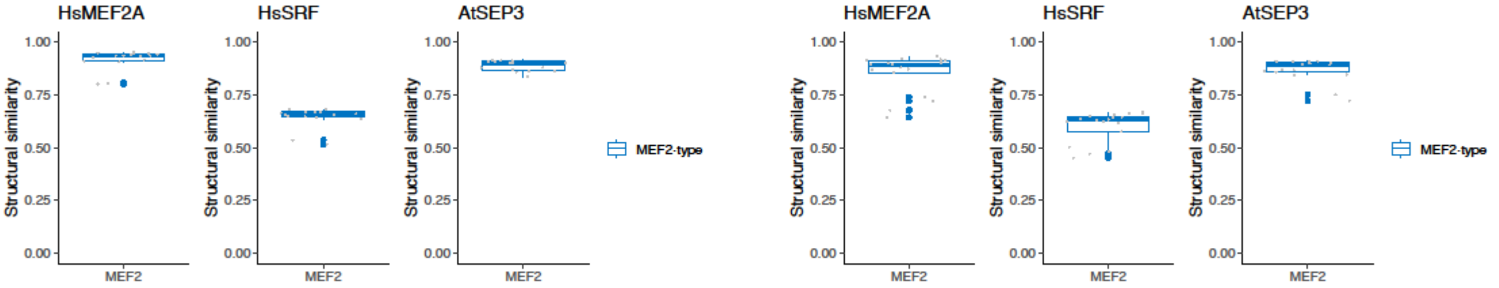


**d.** Land plants

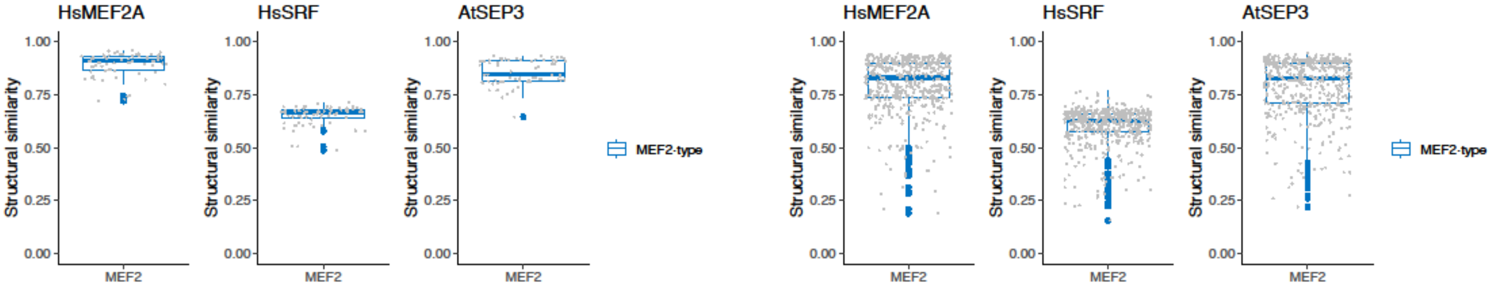

**Fig.S5** Analyses of predicted protein structures of MADS-box transcription factors.

**e.** Predicted structures of plant Type I and II proteins as monomers and dimers by AlphaFold2. Monomers are colored by confidence, red marking high confidence and blue marking low confidence. Dimers are colored to differentiate the two chains. The gray frames locate the helix-strand-helix functional units, the extended definition of MADS domains.

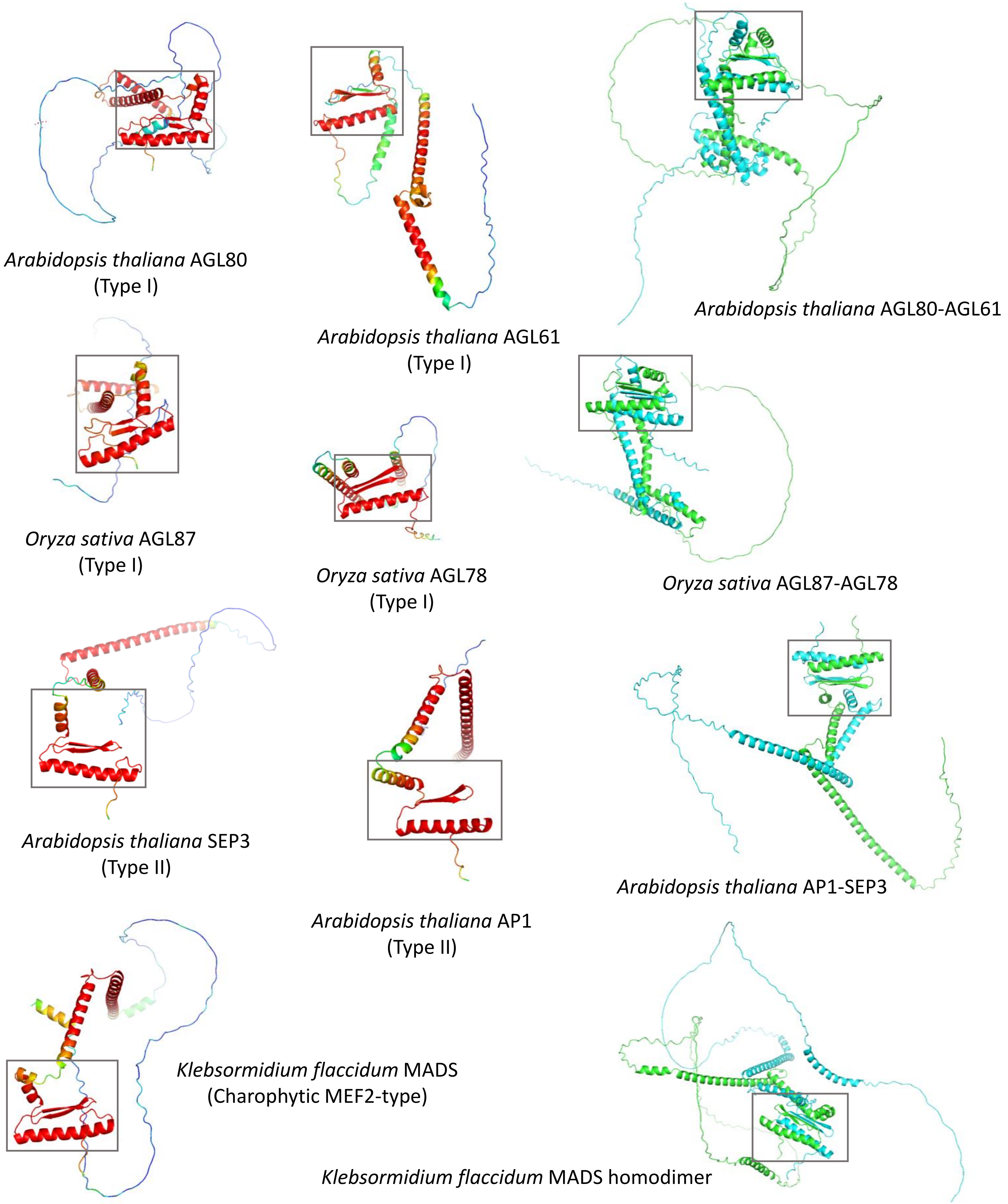

**Fig.S6** Maximum-Likelihood tree of surveyed MADS-box domains with SRF-like sequences in the core Chlorophytes (green clade pointed by the arrow). Bootstrap values are labelled above branches of interest.

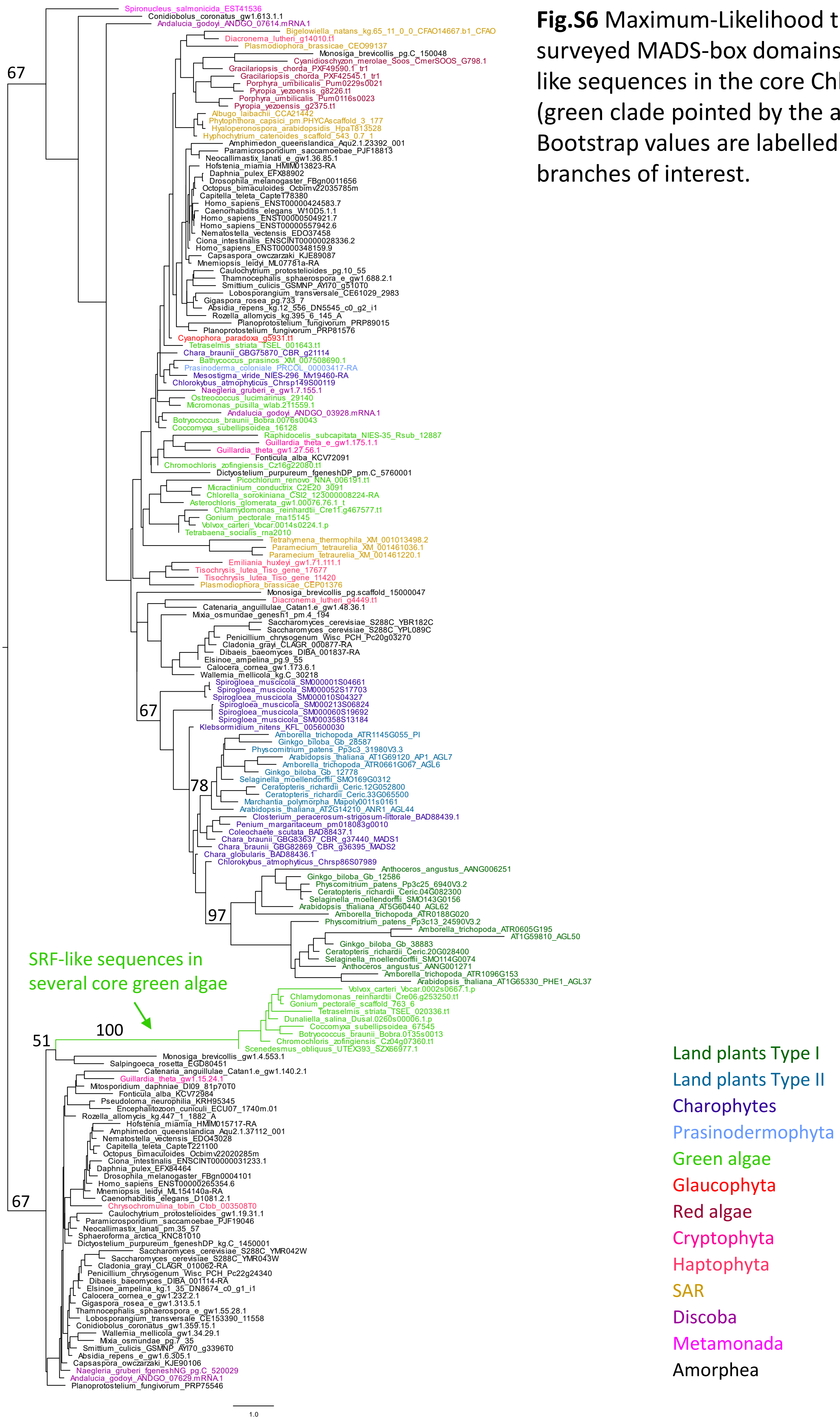

**Fig.S7** Maximum likelihood trees inferred with the alignments generated by two additional softwares, T-Coffee and MAFFT. Extended MADS domain sequences were aligned. The substitution model is LG. Bootstrap values are labelled next to branches of interest.

**a.** T-coffee.

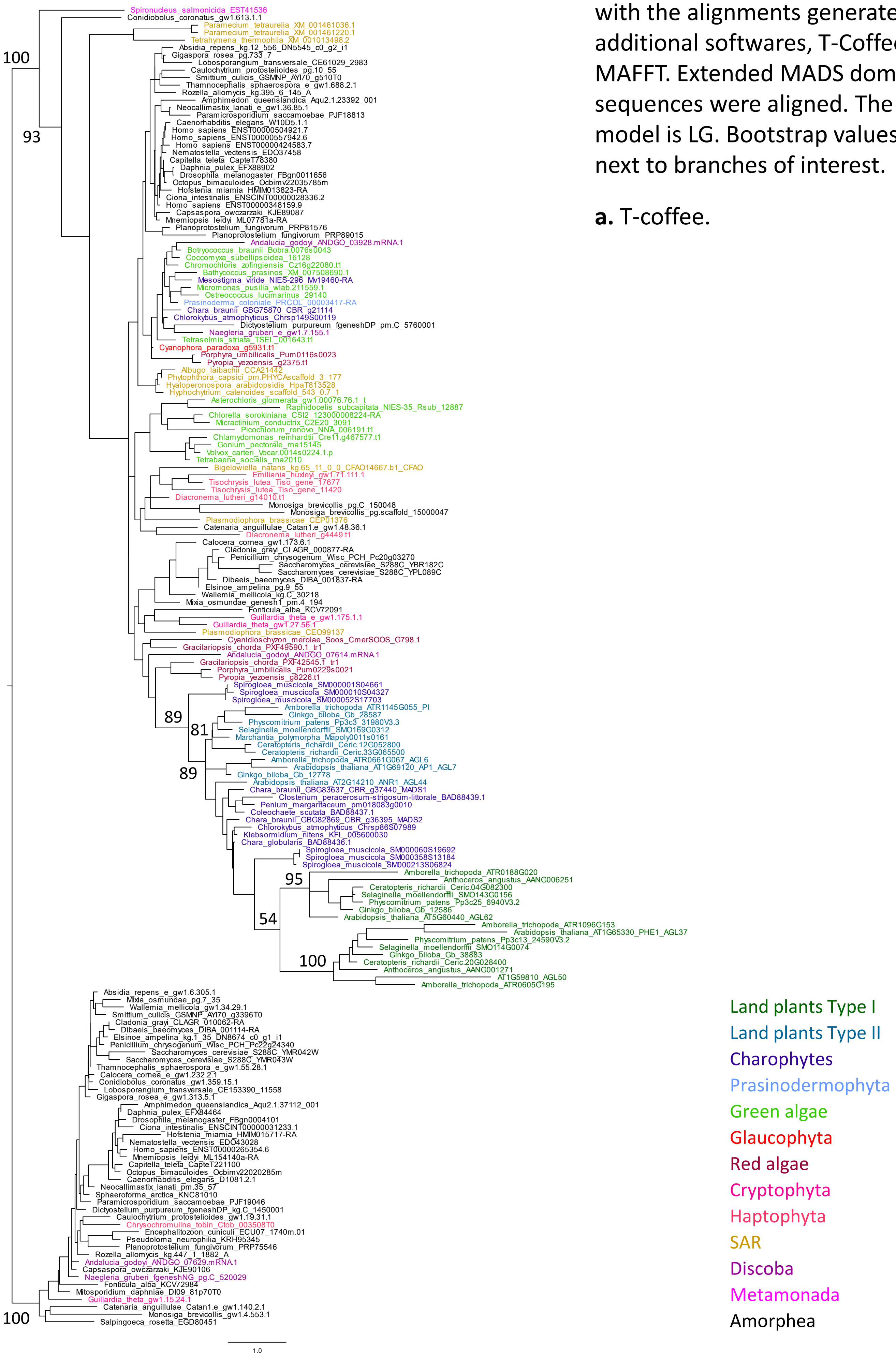

**Fig.S7** Maximum likelihood trees inferred with the alignments generated by two additional softwares, T-Coffee and MAFFT. Extended MADS domain sequences were aligned. The substitution model is LG. Bootstrap values are labelled next to branches of interest.

**b.** MAFFT.

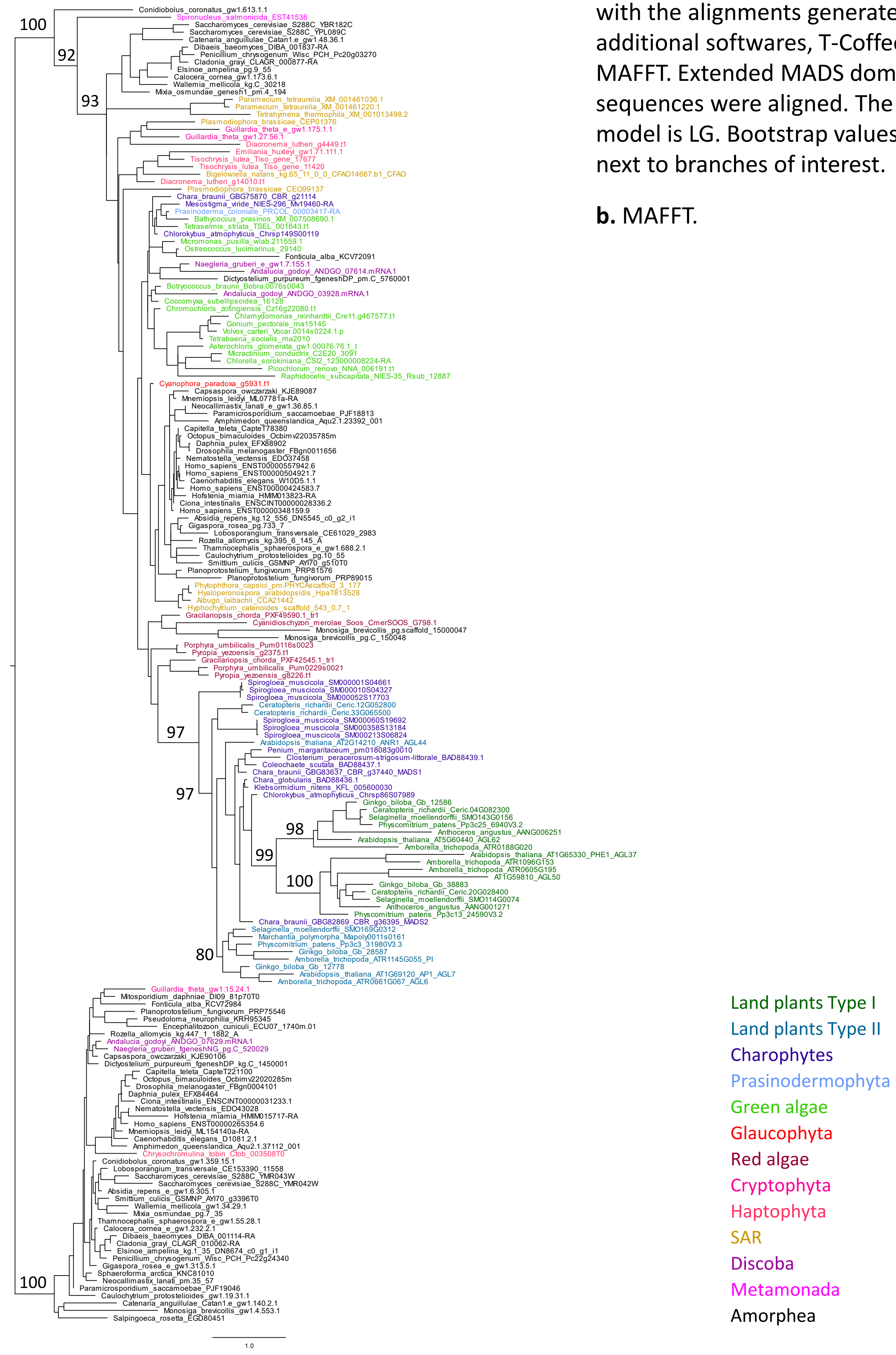

**Fig.S8** Maximum likelihood trees inferred with two additionally substitution models, JTT and WAG. The alignments of extended MADS domain sequences were generated by MUSCLE. Bootstrap values are labelled next to branches of interest.

**a.** JTT.

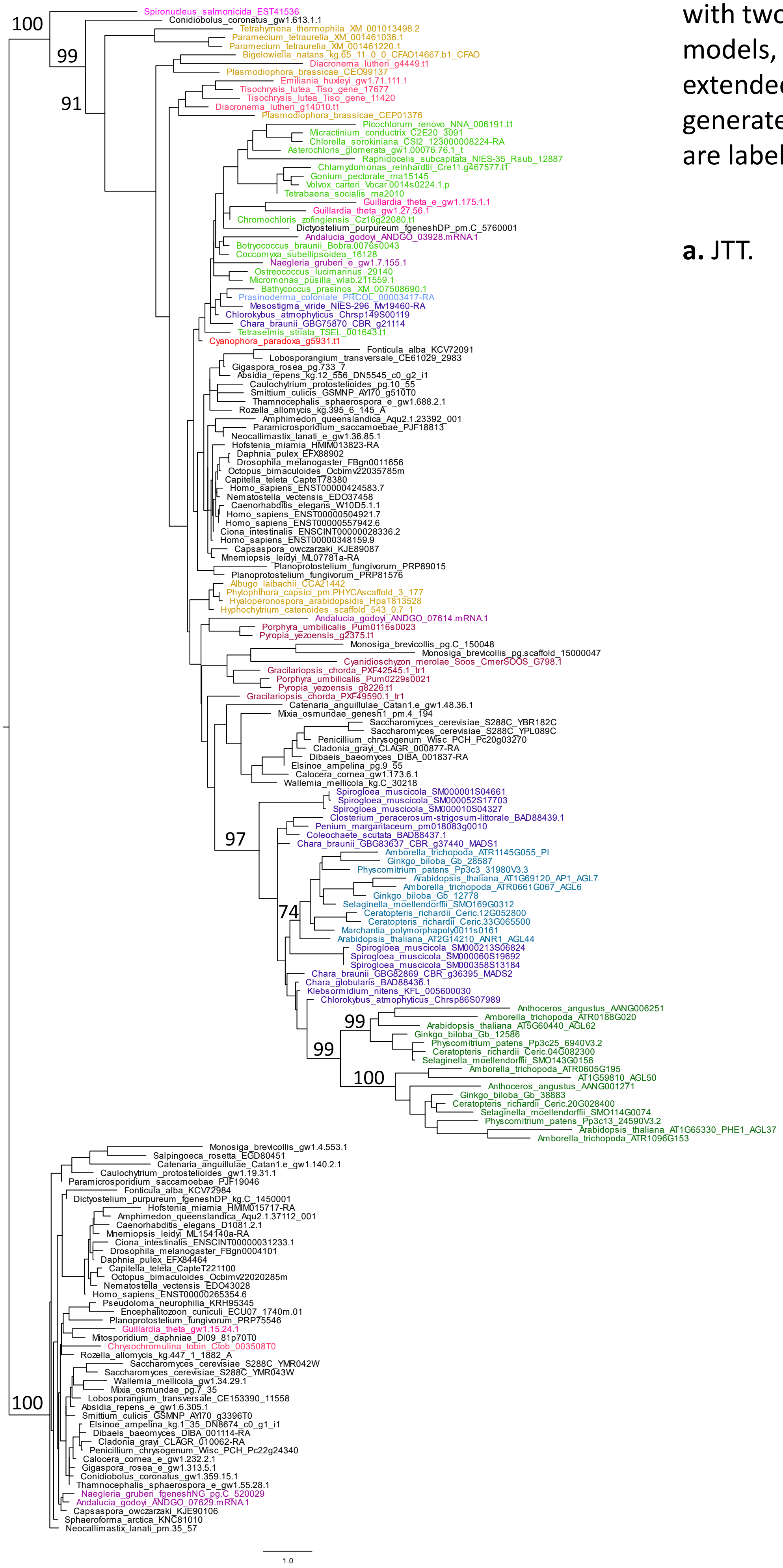


Land plants Type I
Land plants Type II
Charophytes
Prasinodermophyta
Green algae
Glaucophyta
Red algae
Cryptophyta
Haptophyta
SAR
Discoba
Metamonada
Amorphea

**Fig.S8** Maximum likelihood trees inferred with two additionally substitution models, JTT and WAG. The alignments of extended MADS domain sequences were generated by MUSCLE. Bootstrap values are labelled next to branches of interest.

**b.** WAG.

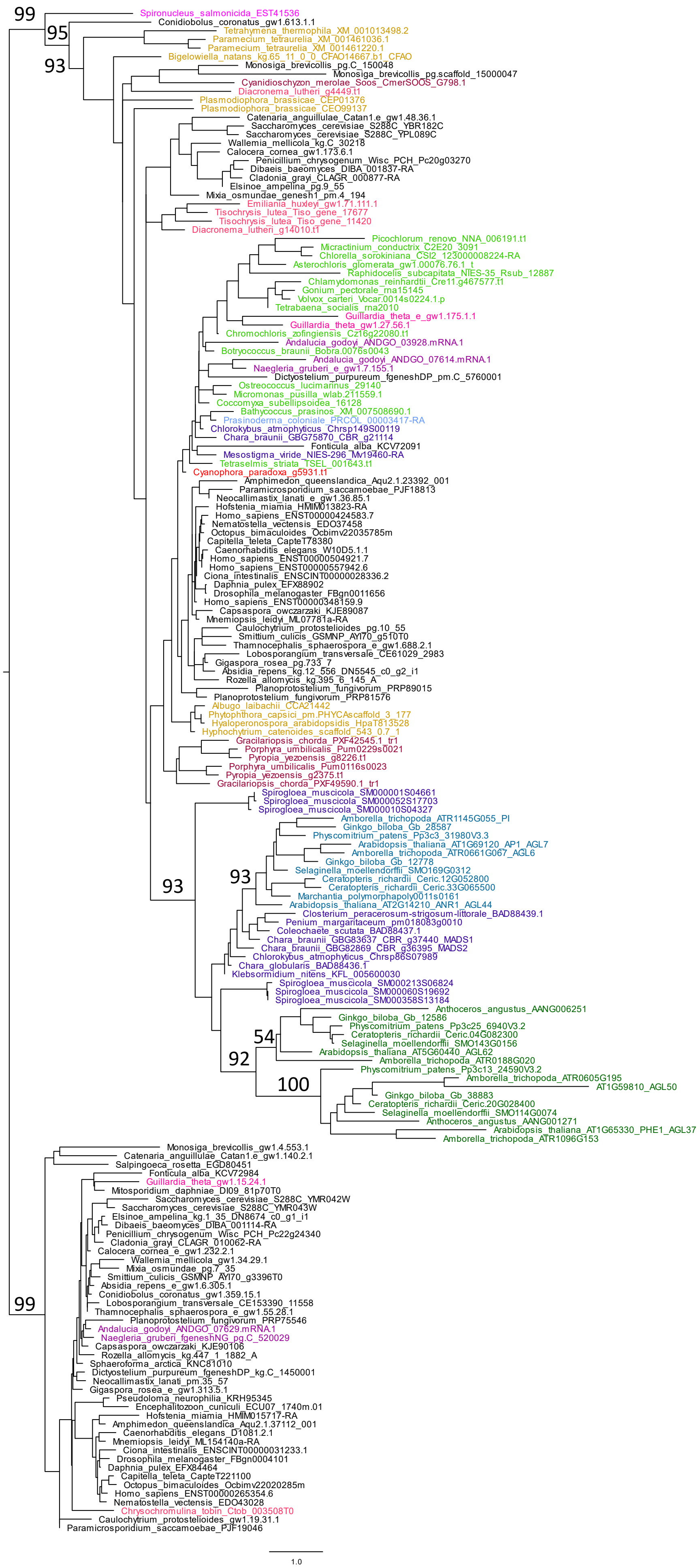


Land plants Type I
Land plants Type II
Charophytes
Prasinodermophyta
Green algae
Glaucophyta
Red algae
Cryptophyta
Haptophyta
SAR
Discoba
Metamonada
Amorphea
